# The Rubicon-WIPI axis regulates exosome biogenesis during aging

**DOI:** 10.1101/2024.05.08.593233

**Authors:** Kyosuke Yanagawa, Akiko Kuma, Maho Hamasaki, Shunbun Kita, Tadashi Yamamuro, Kohei Nishino, Shuhei Nakamura, Hiroko Omori, Tatsuya Kaminishi, Satoshi Oikawa, Yoshio Kato, Ryuya Edahiro, Ryosuke Kawagoe, Takako Taniguchi, Yoko Tanaka, Takayuki Shima, Keisuke Tabata, Miki Iwatani, Nao Bekku, Rikinari Hanayama, Yukinori Okada, Takayuki Akimoto, Hidetaka Kosako, Akiko Takahashi, Iichiro Shimomura, Yasushi Sakata, Tamotsu Yoshimori

**Author notes:** Correspondence to Tamotsu Yoshimori.

## Abstract

Cells release intraluminal vesicles (ILVs) in multivesicular bodies as exosomes to communicate with other cells. Although recent studies suggest an intimate link between exosome biogenesis and autophagy, the detailed mechanism is not fully understood. Here we employed comprehensive RNAi screening for autophagy-related factors and discovered that Rubicon, a negative regulator of autophagy, is essential for exosome release. Rubicon recruits WIPI2d to endosomes to promote exosome biogenesis. Interactome analysis of WIPI2d identified the ESCRT components that are required for ILV formation. Notably, we found that Rubicon is required for an age-dependent increase of exosome release in mice. In addition, small RNA sequencing of serum exosomes revealed that Rubicon determines the fate of exosomal microRNAs associated with cellular senescence and longevity pathways. Taken together, our current results suggest that the Rubicon-WIPI axis functions as a key regulator of exosome biogenesis and is responsible for the age-dependent changes in exosome quantity and quality.

## Main

Exosomes, which are small extracellular vesicles (EVs) with a diameter ranging from 30 to 150 nm, originate from multivesicular bodies (MVBs) and transfer proteins, lipids, and nucleic acids to recipient cells for intercellular communication.^1–7^ Exosome secretion is a complicated multistep process via the endocytic pathway. The intraluminal vesicles (ILVs) bud inward from the endosomal membrane to form MVBs, which fuse with the plasma membrane and release their ILVs as exosomes.^1–4,8^ Indeed, previous reports showed that the endosomal sorting complex required for transport (ESCRT) components and their associated proteins are recruited to the specific sites on the endosome to mediate ILV formation.^9,10^ However, the detailed mechanism of this recruitment in exosome biogenesis remains largely elusive. Furthermore, a recent study showed that the number of circulating exosomes increases with age.^11^ Another report indicated that exosomal contents, including microRNA (miRNA), vary with age.^12,13^ While age-dependent changes in the quantity and quality of exosomes could have a pro-aging effect,^13–17^ the molecular mechanism of exosomal alterations in aging remains unclear.

Macroautophagy (referred to hereinafter as autophagy) is a degradation and recycling process essential for the maintenance of cellular homeostasis. During autophagy, double-membrane organelles known as autophagosomes sequester cytoplasmic components and fuse with lysosomes to degrade their contents. This process is orchestrated by a set of autophagy-related (ATG) genes^18–20^ and is negatively regulated by Run Domain Beclin-1-Interacting And Cysteine-Rich Domain-Containing Protein (Rubicon).^21^ Our previous study suggests that Rubicon accumulates with age, thereby leading to an age-associated decline of autophagy in various organisms.^22^ Other recent studies suggest Rubicon may have multiple autophagy-independent functions.^23–25^ As well as autophagy, MVB formation is a membrane-trafficking pathway involved in the internalization and degradation of cellular components;^26^ therefore, it is plausible that exosome biogenesis and autophagy may share a regulatory mechanism.^27^ Whereas some studies have claimed that autophagy is involved in exosome secretion,^28,29^ interestingly, other studies have suggested that several ATGs, including *ULK1*, *ATG3*, *ATG5*, *ATG7*, *ATG12*, and *ATG16L1*, contribute to exosome release independently of the autophagy pathway.^30–34^ In other words, determining whether exosome biogenesis requires the entire autophagy machinery or some specific autophagy-related factors remains a complex issue.

Here, we report that Rubicon positively regulates exosome biogenesis in an autophagy-independent manner. Rubicon mediates the endosomal recruitment of the WD repeat domain phosphoinositide interacting protein 2 (WIPI2), which interacts with the ESCRT machinery required for MVB formation. Furthermore, we discovered that Rubicon is essential for the age-dependent exosomal secretion of miRNAs associated with cellular senescence and aging-related pathways.

## Results

### RNAi screening identifies Rubicon as a novel regulator of exosome secretion

To determine whether exosome release requires the entire autophagic pathway or only specific autophagy factors, we first conducted RNA interference (RNAi) screening for autophagy-related factors (Extended Data Fig. 1a). We knocked down *ATG*s and *Rubicon* in human mesenchymal stem cells (hMSCs), and isolated exosomes from the culture medium for western blotting. The RNAi screening showed that knockdown of *ULK1*, *ATG5*, or *Rubicon*, but no other genes, decreased the levels of exosome markers, such as CD63 and Flotillin-1, in the exosome fraction (Fig. 1a,b and Extended Data Fig. 1b). Consistent with this, the levels of CD63 and Flotillin-1 in the exosome fraction were reduced in *Atg5-*knockout (KO) mouse embryo fibroblasts (MEFs) (Extended Data Fig. 1c,d), but not in *Fip200-*, *Atg2-*KO or *Atg14*-KO MEFs (Extended Data Fig. 1e–j). The levels of CD63 in the exosome fraction was also reduced in *Atg16l1*-KO MEFs as well as *Atg5*-KO MEFs, and the decrease was rescued by ATG16L1 overexpression (Extended Data Fig. 1k,l). Knockdown of ATG13 significantly increased exosome secretion, however this result was inconsistent with knockdown of ULK1 or FIP200 (Fig. 1a), which are components of the ULK1 complex similar with ATG13. These results suggest that exosome secretion does not require the whole autophagic pathway, but only certain autophagy factors. Importantly, the levels of CD63, ALIX, and Flotillin-1 in exosomes were reduced in *Rubicon-*KO MEFs (Fig. 1c,d), and these levels were rescued by Rubicon expression (Extended Data Fig. 1m). Conversely, the exosomal markers were increased in *Rubicon*-overexpressing (OE) MEFs (Fig. 1e,f, and Extended Data Fig. 1n). Both *RUBCN* knockdown and *Rubicon* OE did not affect cell viability, and, importantly, the levels of exosome markers in the exosome fraction normalized by the number of live cells were reduced in *RUBCN*-knockdown cells (Extended Data Fig. 1o-s). *Rubicon* OE increased the total protein levels in the exosome fraction normalized by the number of live cells (Extended Data Fig. 1r,s). Moreover, the number of small EVs (sEVs), as quantified by nanoparticle tracking analysis (NTA), was reduced in *Rubicon*-KO MEFs (Fig. 1g,h). These findings indicate that the amount of Rubicon is a determinant for exosome secretion. Because previous reports showed that ULK1 and ATG5 are involved in exosome release,^30,31,34^ we sought to clarify the role of Rubicon in exosome secretion. To this end, we assessed the subcellular localization of Rubicon and found that it localized to CD63-positive endosomes (Extended Data Fig. 2a,b). Next, to enlarge endosomes for better visualization, we utilized a FYVE finger-containing phosphoinositide kinase (PIKFyve) inhibitor, Apilimod, which promotes phosphatidylinositol 3–phosphate (PI3P) accumulation and MVB formation (Extended Data Fig. 2c).^35,36^ It is worth noting that multiple puncta of CD63 were observed within the enlarged endosomes whose membrane was positive for both Rubicon and the PI3P probe mCherry-2×FYVE (Fig. 1i and Extended Data Fig. 2d,e). Consistent with previous reports^21^ which show that Rubicon localizes to LAMP1-positive endolysosomes in basal condition, Rubicon also localized to LAMP1-postive endolysosomes under Apilimod treatment (Extended Data Fig. 2f). Importantly, Apilimod treatment failed to increase the levels of CD63 in the exosome fraction in *Rubicon*-KO cells (Extended Data Fig. 2g,h). These results suggest that endosomal PI3P plays a role in MVB formation in a Rubicon-dependent manner.

**Fig. 1.**
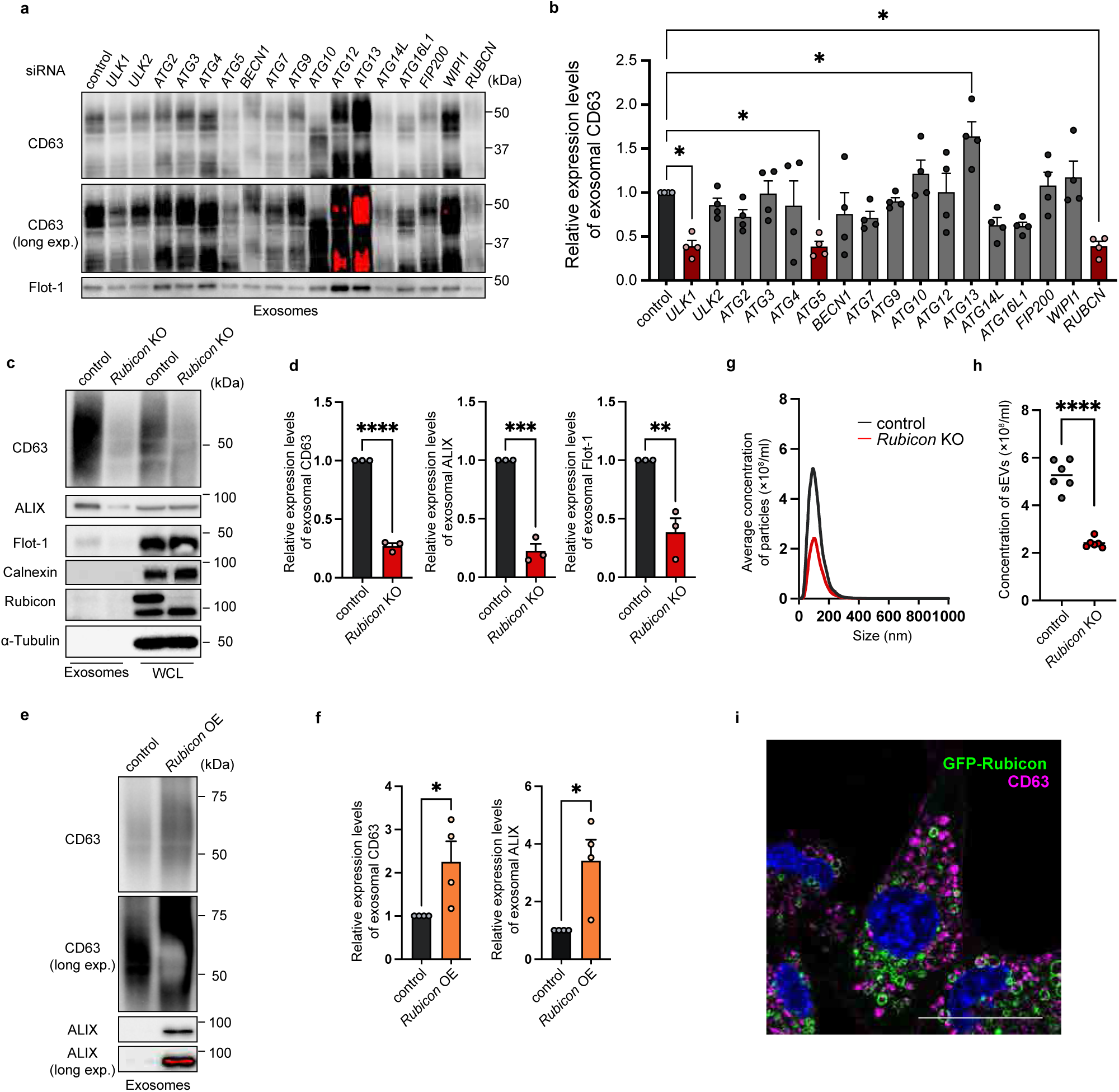
RNAi screening reveals that Rubicon promotes exosome secretion. (a) Immunoblotting of CD63 and Flotilin-1 (Flot-1) in the exosome fractions isolated from the culture medium of human mesenchymal stem cells (hMSCs) with knockdown of the indicated genes. The loading amount for each sample was set as follows: half of the exosome sample collected from 1 ml of culture medium. (b) Quantification of the exosomal CD63 levels in the hMSCs with knockdown of the genes indicated in (a). Bars represent means ± SEM. n = 4 independent experiments. *p < 0.05 (each vs control) by one-way ANOVA followed by Dunnett’s test. (c) Immunoblotting of the indicated proteins in the exosome fractions and the WCLs obtained from control and *Rubicon*-knockout (KO) mouse embryo fibroblasts (MEFs). The loading amount for each sample was set as follows: half of the exosome sample collected from 1 ml of culture medium, and 6 µg for the WCL. (d) Quantification of the exosomal levels of CD63, ALIX, and Flot-1 in (c). Bars represent means ± SEM. n = 3 independent experiments. **p < 0.01, ***p < 0.001, ****p < 0.0001 by a two-tailed Student’s t-test. (e) Immunoblotting of CD63 and ALIX in the exosome fractions isolated from the cultured medium of control and Rubicon-overexpressing (OE) MEFs. The loading amount for each sample was set as follows: half of the exosome sample collected from 1 ml of culture medium. (f) Quantification of the exosomal CD63 and ALIX levels in (e). Bars represent means ± SEM. n = 4 independent experiments. *p < 0.05 by a two-tailed Student’s t-test. (g) Nanoparticle tracking analysis (NTA) of the extracellular vesicles (EVs) purified from the cultured medium of control or *Rubicon*-KO MEFs using a phosphatidylserine affinity magnetic resin (PS-affinity kit). (h) Quantification of the total particle concentration of the EVs in (g). Bars represent means. n = 6 independent experiments. ****p < 0.0001 by a two-tailed Student’s t-test. (i) Immunofluorescent images of CD63 (magenta) and DAPI (blue) in MEFs stably expressing GFP-Rubicon treated with 0.5 μM Apilimod for 1 hour. Scale bars, 50 μm.

### Rubicon regulates exosome secretion independently of both canonical and non-canonical autophagy

Previous studies have shown that Rubicon forms a complex with UVRAG and Beclin-1 to regulate autophagy and its non-canonical forms, such as LC3-associated phagocytosis (LAP) and LC3-associated endocytosis (LANDO).^21,23,24^ Therefore, we examined whether UVRAG or Beclin-1 is also required for exosome release and found that both UVRAG and Beclin-1 were dispensable for exosome secretion (Extended Data Fig. 3a–c) and localization of Rubicon to MVBs (Extended Data Fig. 2a,b,d). A previous study suggests that Rubicon forms a complex independently of Beclin-1, based on co-fractionation data.^37^ However, similar experiments conducted in our previous study did not yield comparable results, likely due to a difference in focus on the complex size.^21^ To clarify this discrepancy, we performed gel filtration experiments more suitable for separating protein complexes smaller than the Beclin-1-containing complex, and successfully eluted fractions containing Rubicon, fraction 25-29, and PIK3C3, fraction 23-29, but not Beclin-1 (Extended data Fig. 3d). Moreover, the WD repeat domain of ATG16L1, which is required for LC3 lipidation in LAP or LANDO, was also dispensable for exosome secretion (Extended Data Fig. 3e,f). These findings support that Rubicon, but not Beclin-1 is not, is necessary for exosome secretion through a previously unknown pathway.

### WIPI2 and WIPI3 interact with Rubicon and promote exosome secretion

To investigate how Rubicon regulates exosome secretion, we sought to identify its interacting partner during exosome biogenesis. Using immunoprecipitation and mass spectrometry, we identified 79 potential interactors of Rubicon in HeLa Kyoto cells (Extended Data Table. 1). We conducted RNAi screen (Extended Data Fig. 1a) of the 79 candidates, and then identified 16 genes whose knockdown decreased the levels of exosomal CD63 to less than a quarter compared with control (Fig. 2a and Extended Data Fig. 4a). Despite our findings that endosomal PI3P plays a pivotal role in exosome secretion (Extended Data Fig. 2c,e,g,h), the 16 candidate genes did not contain PIK3C3 and PIK3R4. Therefore, we sought to determine the factor responsible for generating the endosomal PI3P necessary for exosome formation. Our further analysis revealed that siPIK3C3 #2, which exhibited a higher knockdown efficacy than siPIK3C3 #1 used in the RNAi screening, significantly decreased exosome markers in the exosome fraction (Extended Data Fig. 4d-f), suggesting that PIK3C3 is essential for exosome secretion. To narrow down the candidates, we used Apilimod to promote exosome biogenesis and examined the interactome of Rubicon in MEFs with Apilimod treatment. Among the 16 identified candidates, we discovered that WIPI2 was strongly co-immunoprecipitated with Rubicon under Apilimod treatment (Fig. 2b and Extended Data Fig. 4b,c; Extended Data Table. 2). Consistent with the RNAi screening results, knockdown or knockout of *WIPI2/Wipi2* impaired exosome release in both western blots and NTA (Fig. 2c–h), suggesting that WIPI2 is required for exosome secretion. WIPI2 is a member of the WIPI family proteins, which are mammalian orthologues of yeast Atg18. The WIPI family is classified as a β-propeller that binds to polyphosphoinositides (PROPPINs) and thus functions as a PI3P-binding effector essential for autophagosome formation.^38–40^ While the function of the WIPI family in the autophagic pathway is well examined, their role in exosome secretion remains largely unexplored. Therefore, we examined whether other WIPI family proteins are required for exosome secretion. WIPI3, like WIPI2, interacted with Rubicon, and this interaction was enhanced under Apilimod treatment (Fig. 2b). Consistent with these findings, exosome release was not reduced in *WIPI1*- or *WIPI4*-deficient cells but was reduced in *WIPI2*- or *WIPI3*-deficient cells (Extended Data Fig. 5a-d). These results suggest that the WIPI proteins, WIPI2 and WIPI3, interact with Rubicon and promote exosome secretion.

**Fig. 2.**
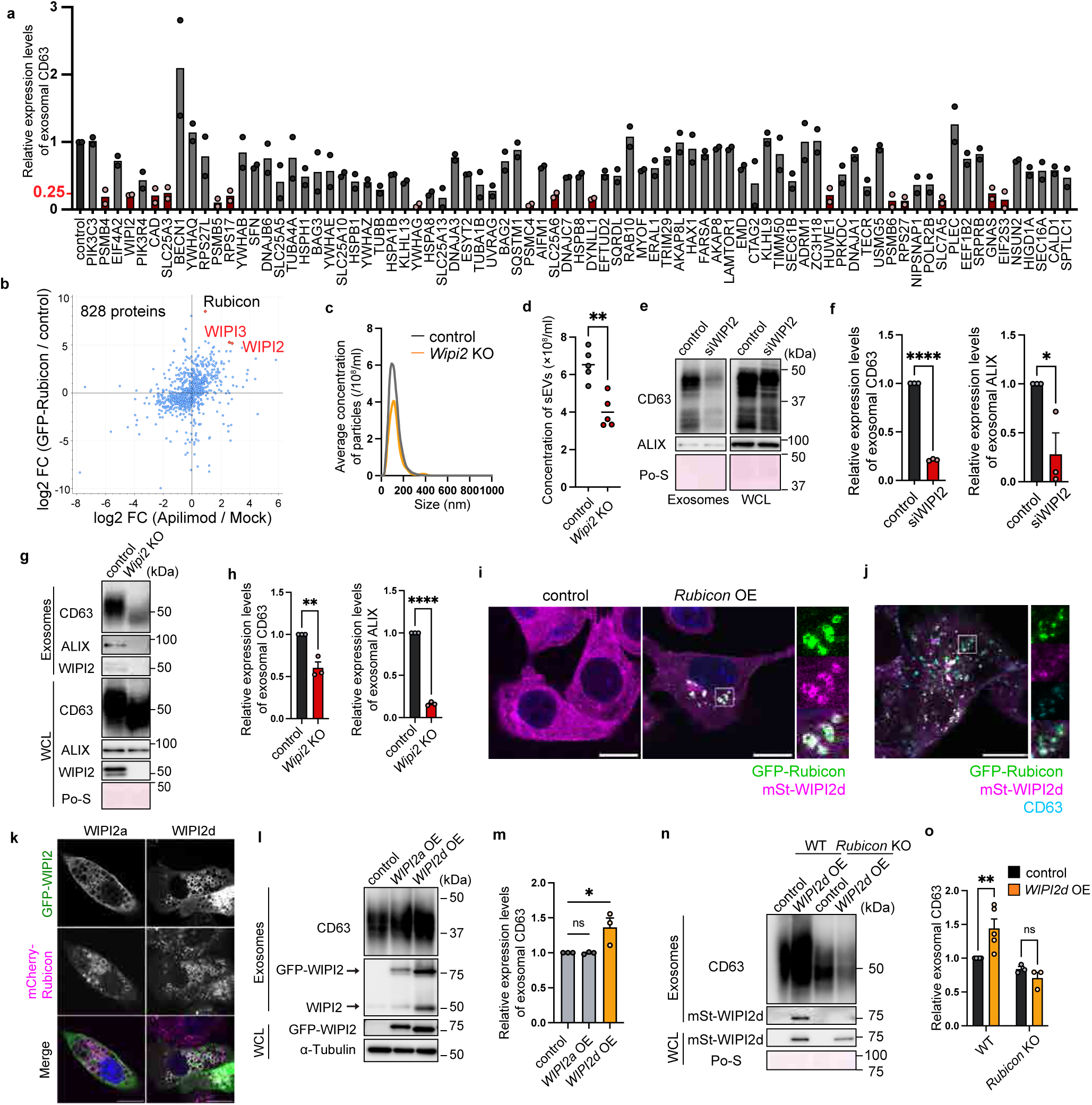
WIPI2 interacts with Rubicon and is required for exosome secretion. (a) Quantification of exosomal CD63 levels in control or knockdown hMSCs with the genes indicated in Fig. S4a. Bars represent means. n = 2 independent experiments. The loading amount for each sample was set as follows: half of the exosome sample collected from 1 ml of culture medium. (b) Label-free quantification of Rubicon interactors. Dots indicate 828 proteins identified and quantified in the LC-MS/MS analysis of the cell immunoprecipitates using anti-GFP nanobody. Control or GFP-Rubicon expressing MEFs were treated with or without 0.5 µM Apilimod for 1 hour. X-axis, log_2_ fold change (FC) between GFP-Rubicon expressing cells treated with or without Apilimod (Apilimod/Mock); Y-axis, log_2_ FC between GFP-Rubicon–expressing and control cells (GFP-Rubicon/control) treated with Apilimod. (c) NTA tracing of the EVs purified from the cultured medium of the indicated cells. (d) Quantification of the total particle concentration of the EVs in (c). Bars represent means. n = 5 independent experiments. **p < 0.01 by a two-tailed Student’s t-test. (e) Immunoblotting of the indicated proteins in the exosome fractions and WCLs obtained from control and *WIPI2* knockdown hMSCs. The loading amount for each sample was set as follows: half of the exosome sample collected from 1 ml of culture medium, and 6 µg for the WCL. (f) Quantification of exosomal CD63 and ALIX levels in (e). Bars represent means ± SEM. n = 3 independent experiments. *p < 0.05, ****p < 0.0001 by a two-tailed Student’s t-test. (g) Immunoblotting of CD63 and ALIX in the exosome fractions and WCLs obtained from control and *WIPI2*-KO MEFs. The loading amount for each sample was set as follows: half of the exosome sample collected from 1 ml of culture medium, and 6 µg for the WCL. (h) Quantification of exosomal CD63 and ALIX levels in (g). Bars represent means ± SEM. n = 3 independent experiments. **p < 0.01, ****p < 0.0001 by a two-tailed Student’s t-test. (i,j) Immunofluorescence images of CD63 (cyan) and DAPI (blue) in MEFs stably expressing mStrawberry-WIPI2d (mSt-WIPI2d) with or without stable expression of GFP-Rubicon. Scale bars, 50 μm. (k) Immunofluorescence images of DAPI (blue) in MEFs stably expressing mCherry-Rubicon with stable expression of GFP-WIPI2a or GFP-WIPI2d. Scale bars, 50 μm. (l) Immunoblotting of the indicated proteins in the exosome fractions and WCLs obtained from control and GFP-WIPI2 stably overexpressing MEFs. The loading amount for each sample was set as follows: half of the exosome sample collected from 1 ml of culture medium, and 6 µg for the WCL. (m) Quantification of the exosomal CD63 levels in the MEFs with overexpressing of the genes indicated in (l). Bars represent means ± SEM. n = 3 independent experiments. ns, not significant; *p < 0.05 (each vs control) by one-way ANOVA followed by Dunnett’s test. (n) Immunoblotting of the indicated proteins in the exosome fractions and WCLs obtained from control and *Rubicon*-KO MEFs with or without stable expression of WIPI2d. The loading amount for each sample was set as follows: half of the exosome sample collected from 1 ml of culture medium, and 6 µg for the WCL. (o) Quantification of exosomal CD63 levels in (n). Bars represent means ± SEM. Wild type (WT), n = 5; *Rubicon* KO, n = 3. ns, not significant; **p < 0.01 by two-way ANOVA.

### Rubicon recruits WIPI2d and WIPI3 to endosomes to promote exosome secretion

Next, we sought to investigate how WIPI2 regulates exosome secretion in concert with Rubicon. Since WIPI2 contains multiple isoforms,^38^ we tested which WIPI2 isoform strongly interacts with Rubicon, and found that WIPI2d was the strongest interactor (Extended Data Fig. 5e). WIPI2d was localized to the cytosol in control cells, but to CD63-positive endosomes in *Rubicon*-OE cells (Fig. 2i,j), suggesting that Rubicon recruits WIPI2 to endosomes.

Additionally, unlike WIPI1, endogenous WIPI2 and WIPI3 localized to Rubicon-decorated endosomes under Apilimod treatment (Extended Data Fig. 5f,g). Endosomal recruitment of WIPI2 was observed in *Fip200*-deficient cells (Extended Data Fig. 5f), demonstrating that it is independent of autophagosome-related pathways such as amphisome formation^27,41,42^ or autophagic lysosome reformation.^43–46^ In WIPI2 isoforms, WIPI2d, but not WIPI2a, was recruited to Rubicon-decorated endosomes (Fig. 2k). Consistent with this, the overexpression of WIPI2d, but not WIPI2a, increased the exosomal levels of CD63 and WIPI2 (Fig. 2l,m). These results suggest that only the WIPI proteins specifically recruited to Rubicon-decorated endosomes promote exosome biogenesis. Importantly, increased exosome secretion upon WIPI2d overexpression was not observed in the absence of Rubicon (Fig. 2n,o, and Extended Data Fig. 5h). This indicates that Rubicon recruits specific WIPI proteins to endosomes in order to promote exosome secretion.

To determine the detailed mechanism of this recruitment, we generated a series of mutant Rubicon proteins, each of which lacked one of the following: an N-terminal serine-rich region (SR-N), a coiled-coil domain (CCD), a C-terminal serine-rich region (SR-C), a helix-coil-rich region (HCR), or a C-terminal homology domain (RHD) (Extended Data Fig. 6a). The ΔC (C terminus–lacking) and the ΔHCR (HCR-lacking) mutant, but no other mutants, failed to pull down endogenous WIPI2 and HA-WIPI2d (Extended Data Fig. 6b,c). Consistent with this, expression of the ΔC mutant Rubicon did not rescue the defect of exosomal markers in the exosome fraction (Extended Data Fig. 6d,e). The ΔHCR mutant was not immunoprecipitated with HA-WIPI2d (Extended Data Fig. 6f). In addition, WIPI3 was pulled down with wild-type Rubicon but not the ΔC mutant and this co-immunoprecipitation was enhanced under Apilimod treatment (Extended Data Fig. 6g). These findings suggest that Rubicon interacts with WIPI proteins through its HCR.

### The Rubicon-WIPI axis is required for MVB formation

We next aimed to determine how the Rubicon-WIPI axis regulates exosome secretion. Because bafilomycin A1 saturates exosome secretion by inhibiting MVB acidification,^47,48^ we used it to assess the maximum exosome release, which reflects exosome production. Bafilomycin A1 increased exosome secretion in wild-type cells, but did not increase it in *Rubicon*-, *WIPI2-* or *WIPI3*-deficient cells (Fig. 3a,b and Extended Data Fig. 7a,b). Double knockdown of *WIPI2* and *WIPI3* suppressed exosome secretion further as compared to single knockdown of either *WIPI2* or *WIPI3* (Extended Data Fig. 7a,b). These findings suggest that exosome secretion requires the Rubicon-WIPI axis, in which WIPI2 and WIPI3 have a redundancy. Therefore, we examined MVB biogenesis using cells expressing Rab5Q79L, a constitutively active mutant of Rab5. Rab5Q79L disrupts endosomal trafficking and promotes the formation of enlarged endosomes containing multiple ILVs.^49–51^ The number of CD63-positive ILVs within the enlarged endosomes was significantly reduced in *WIPI2* knockdown cells (Fig. 3c,d). As described above, Apilimod also promotes MVB formation. Under Apilimod treatment, the CD63-positive ILVs surrounded by Rubicon-positive endosomes were reduced in number in *WIPI2*-deficient cells (Fig. 3e,f). Electron microscopy showed a decrease in the number of MVBs in *Rubicon*-deficient cells (Extended Data Fig. 7c,d). Consistently, immunoelectron microscopy revealed that the number of MVBs, which contains GFP-CD63-decorated ILVs, was reduced in *Rubicon*-deficient cells (Fig. 3g,h and Extended Data Fig.7e). Taken together, our results demonstrate that the Rubicon-WIPI axis is essential for MVB formation.

**Fig. 3.**
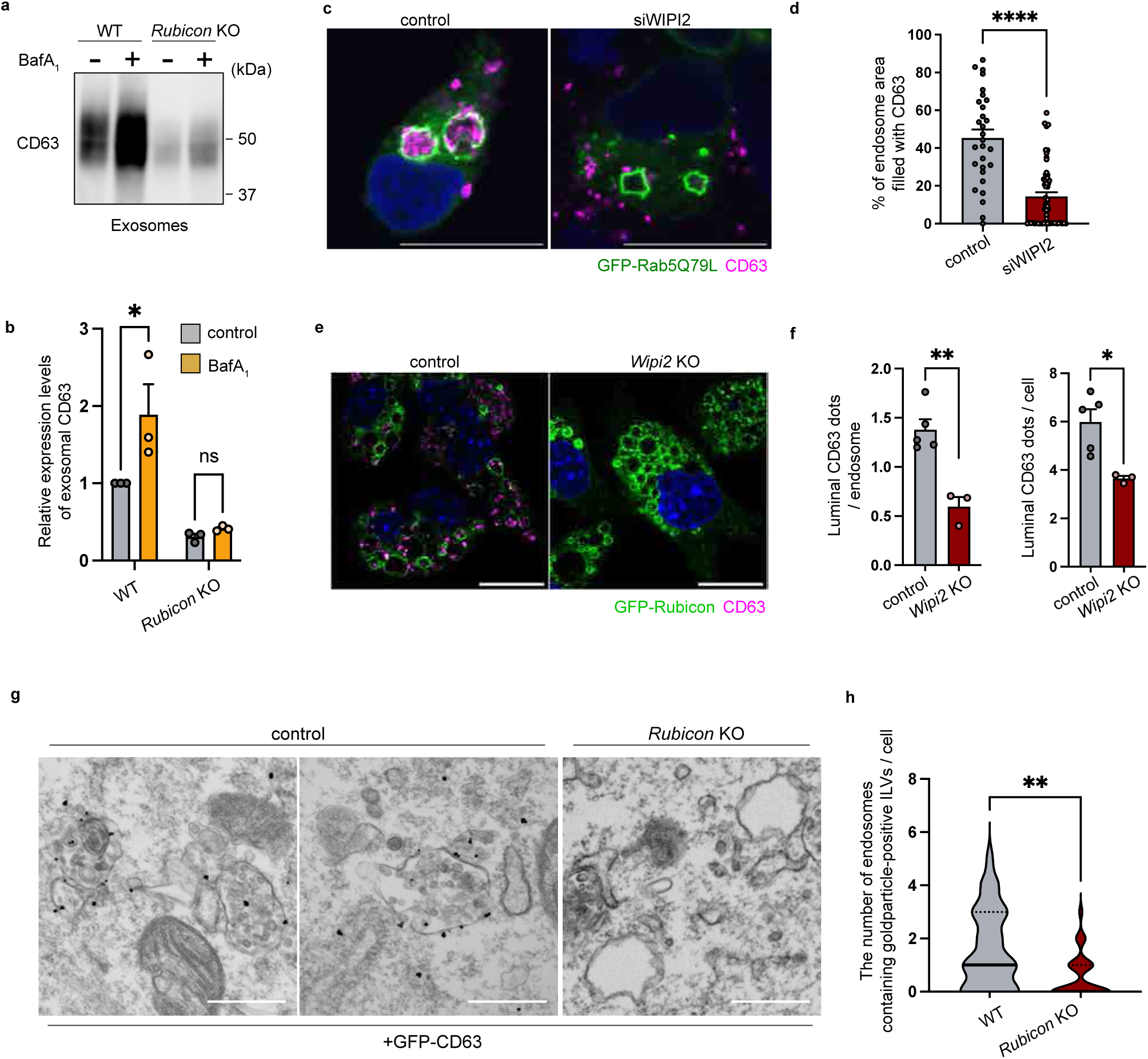
The Rubicon-WIPI axis is required for MVB formation. (a) Immunoblotting of CD63 in the exosome fractions obtained from control and *Rubicon*-KO MEFs with or without bafilomycin A1 treatment for 3 hours. The loading amount for each sample was set as follows: half of the exosome sample collected from 1 ml of culture medium. (b) Quantification of exosomal CD63 levels in (a). Bars represent means ± SEM. n = 3 independent experiments. ns, not significant; *p < 0.05 by two-way ANOVA. (c) Immunofluorescent images of CD63 (magenta) and DAPI (blue) in *WIPI2* (siWIPI2) or *Luciferase* (control) knockdown HEK293T cells transfected with GFP-Rab5Q79L. Scale bars, 50 μm. (d) Quantification of the filling rate of GFP-Rab5Q79L–decorated vacuoles with CD63 in (c). n = 20. ****p < 0.0001 by a two-tailed Student’s t-test. (e) Immunofluorescence images of CD63 (magenta) and DAPI (blue) in control and *WIPI2*-deficient MEFs stably expressing GFP-Rubicon with Apilimod treatment. Scale bars, 50 μm. (f) Quantification of the number of CD63 dots in the GFP-Rubicon–decorated vacuoles in (e). n = 5 independent analysis with more than 20 cells in each condition; *p < 0.05, **p < 0.01 by a two-tailed Student’s t-test. (g) Representative immunoelectron microscopic images of multivesicular bodies (MVBs) in control and *Rubicon*-KO MEFs stably expressing GFP-CD63. (h) Violin plots showing the number of MVBs per cell in (g). WT, n = 33; *Rubicon* KO, n = 31. The solid line denotes the median, and dotted lines define the quartiles; **p < 0.01 by a two-tailed Student’s t-test.

### WIPI2 interacts with multiple ESCRT components and promotes the optimal recruitment of ESCRT machinery in a Rubicon-dependent manner

To obtain further insight into the regulation of MVB formation via the Rubicon-WIPI axis, we performed interactome analysis of WIPI2d and identified multiple ESCRT and associated proteins, including HRS, ALIX, VPS4B, VTA1, and IST1 (Fig. 4a,b, and Extended Data Fig. 8a and Table 3). Since there are several discrepancies between studies regarding which ESCRT factors control exosome biogenesis,^10,50–56^ we knocked down ESCRT factors and found that knockdown of *HRS*, *ALIX*, and *VPS4* decreased exosomal markers (Extended Data Fig. 8b) Knockdown of *HRS*, *ALIX*, *TSG101*,and *VPS4* decreased the particle number of sEVs in NTA in hMSCs (Extended Data Fig. 8c). These results suggest that WIPI2d-interacting ESCRT proteins are essential for exosome formation. Because the accumulation of ESCRT components on MVBs was not observed in basal conditions (data not shown), we employed a dominant-negative VPS4 (VPS4-EQ), which impairs disassembly of ESCRT components and makes it possible to visualize ESCRT localization to MVBs.^57^ The VPS4-EQ–positive MVBs were reduced in *Rubicon*-KO cells (Fig. 4c,d, and Extended Data Fig. 8d). We also found that *RUBCN* KO decreased ALIX accumulation on endosomes in *VPS4*-knockdown cells (Fig. 4e,f), previously used for evaluating ESCRT recruitment.^58^ These observations suggest that Rubicon is necessary for ESCRT recruitment to endosomes. Because ESCRT-0, HRS, recruits the downstream ESCRT factors, we examined whether Rubicon is required for recruitment of HRS onto endosomes^56^. Knockdown of *RUBCN* decreased the interaction of HRS with EGFR, a substrate on endosomes for ILV formation (Extended data Fig. 8e), suggesting that the Rubicon-WIPI axis is necessary for the recruitment of HRS to endosomes. Importantly, GFP-WIPI2d pulled down an ESCRT component, HRS, in a Rubicon-dependent manner (Fig. 4g), whereas ESCRT proteins were not co-immunoprecipitated with Rubicon (Extended data Fig. 8f). Rubicon was not detected in the exosome fraction in western blotting (Fig. 1c). Taken together, our results suggest that Rubicon promotes the recruitment of WIPI2 to endosomes, where WIPI2 recruits ESCRT proteins to endosomes for ILV formation (Fig. 4h).

**Fig. 4.**
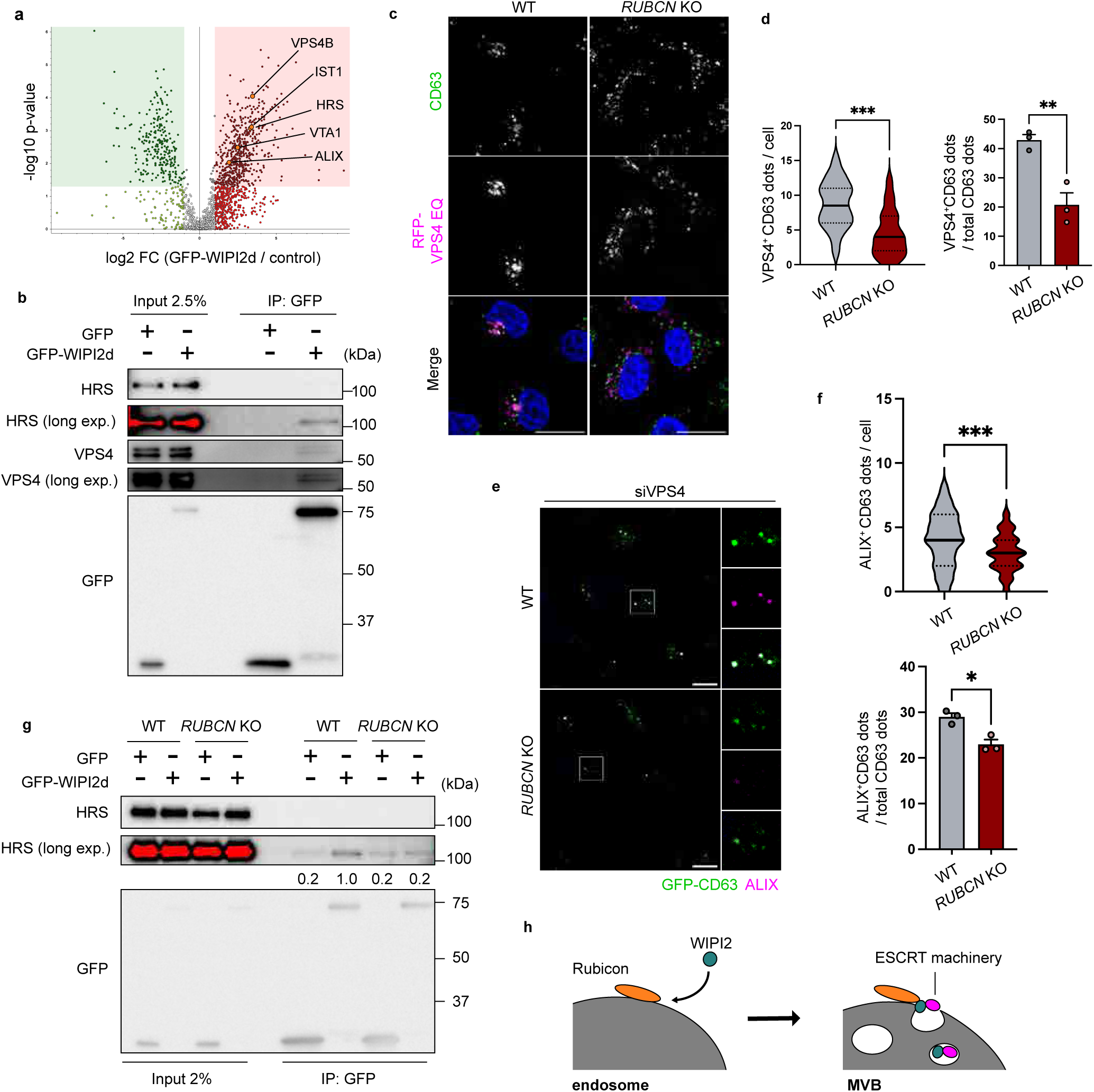
WIPI2 interacts with multiple ESCRT components in a Rubicon-dependent manner. (a) Volcano plot showing label-free quantification of the WIPI2d interactors. Dots indicate proteins identified and quantified in the LC-MS/MS analysis of the immunoprecipitates from control and GFP-WIPI2d–expressing MEFs. Immunoprecipitation was conducted using anti-GFP nanobodies. X-axis, log_2_ FC of GFP-WIPI2d–expressing vs control cells; Y-axis, −log_10_ p-value of triplicate (n = 3) experiments. The red area shows significantly increased proteins (FC > 2, p < 0.05). The green area shows significantly decreased proteins (FC < 2, p < 0.05). (b) Immunoprecipitation assay. HEK293T cells were transfected with the indicated plasmids for 24 hours, and then the cells were lysed and immunoprecipitated with anti-GFP nanobodies. Precipitates were subjected to immunoblotting with the indicated antibodies. n = 3 independent experiments with similar results. (c) Immunofluorescence images of CD63 (green) and DAPI (blue) in HeLa Kyoto cells transfected with mRFP-VPS4E228Q (RFP-VPS4EQ). Scale bars, 50 μm. (d) Left: Violin plot showing the number of mRFP-VPS4E228Q–positive endosomes (Green^+^ Magenta^+^ dots) in the indicated cells. n = 21, The solid line denotes the median, and the dotted lines define the quartiles; ***p < 0.001 by a two-tailed Student’s t-test. Right: Quantification of mRFP-VPS4E228Q–positive endosomes per total endosomes in the indicated cells. n = 3 independent analyses with more than 20 cells in each condition; **p < 0.01 by a two-tailed Student’s t-test. (e) Immunofluorescence images of ALIX (magenta) and DAPI (blue) in control and *RUBCN*-deficient HeLa Kyoto cells transfected with siVPS4 and GFP-CD63. Scale bars, 50 μm. (f) Upper: Violin plot showing the number of ALIX- and CD63-positive endosomes (Green^+^ Magenta^+^ dots) in the indicated cells. WT, n = 76; *Rubicon* KO, n = 72., The solid line denotes the median, and the dotted lines define the quartiles; ***p < 0.001 by a two-tailed Student’s t-test. Bottom: Quantification of ALIX- and CD63-positive endosomes (Green^+^ Magenta^+^ dots) per total endosomes in the indicated cells. n = 3 independent analyses with more than 20 cells in each condition; *p < 0.05 by a two-tailed Student’s t-test. (g) Immunoprecipitation assay. WT or *Rubicon*-KO HeLa cells were transfected with the indicated plasmids for 24 hours, and then the cells were lysed and immunoprecipitated with anti-GFP nanobodies. Precipitates were subjected to immunoblotting with the indicated antibodies. n = 3 independent experiments with similar results. (h) Illustration of Rubicon, WIPI2, and ESCRT machinery in the MVB formation process.

### Rubicon is necessary for the age-dependent increase in exosome secretion in mice

To examine the role of the Rubicon-WIPI axis *in vivo*, we assessed exosome secretion in *Rubicon*-KO mice (Fig. 5a), which did not exhibit any growth defect that could affect exosome secretion (Extended Data Fig. 9a,b). We isolated serum small EVs^59^ (Extended Data Fig. 9c) and found that *Rubicon*-KO mice showed a significantly decreased number of serum small EVs (Fig. 5b,c and Extended Data Fig. 9d).). On the other hand, *Rubicon*-KO mice did not exhibit a decrease in the number of serum large EVs (Extended Data Fig. 9e). Adipose tissue–derived mesenchymal stem cells (AD-MSCs), which are a major exosome source, exhibited reduced exosome production in *Rubicon*-KO mice (Fig. 5d,e and Extended Data Fig. 9f). Of note, our recent study showed that Rubicon accumulates with age across several tissues and species;^22^ thus, we hypothesized that its accumulation would promote exosome release in aged organisms. As expected, Rubicon was significantly increased in AD-MSCs isolated from aged mice (Extended Data Fig. 9g,h). Consistent with this, circulating exosomes were increased in aged wild-type mice, and strikingly, this increase was suppressed in aged *Rubicon*-KO mice (Fig. 5f-i and Extended Data Fig. 9i). These results suggest that Rubicon is essential for the age-dependent increase in exosome secretion in mice. Previous reports showed that cellular senescence promotes secretion of small EVs^60^ and also increases the expression of Rubicon^61^, therefore, we further investigated the role of Rubicon in exosome secretion during cellular senescence. Consistent with the previous reports, either of doxorubicin (DXR) -induced senescence or replicative senescence significantly increased exosome secretion (Fig. 5j,k and Extended Data Fig. 9j). This increase was suppressed by *Rubicon* KO (Fig. 5j,k), suggesting that Rubicon play a vital role in exosome secretion associated with cellular senescence. Taken together, our findings suggest that Rubicon accumulates with age and promotes exosome secretion in mice.

**Fig. 5.**
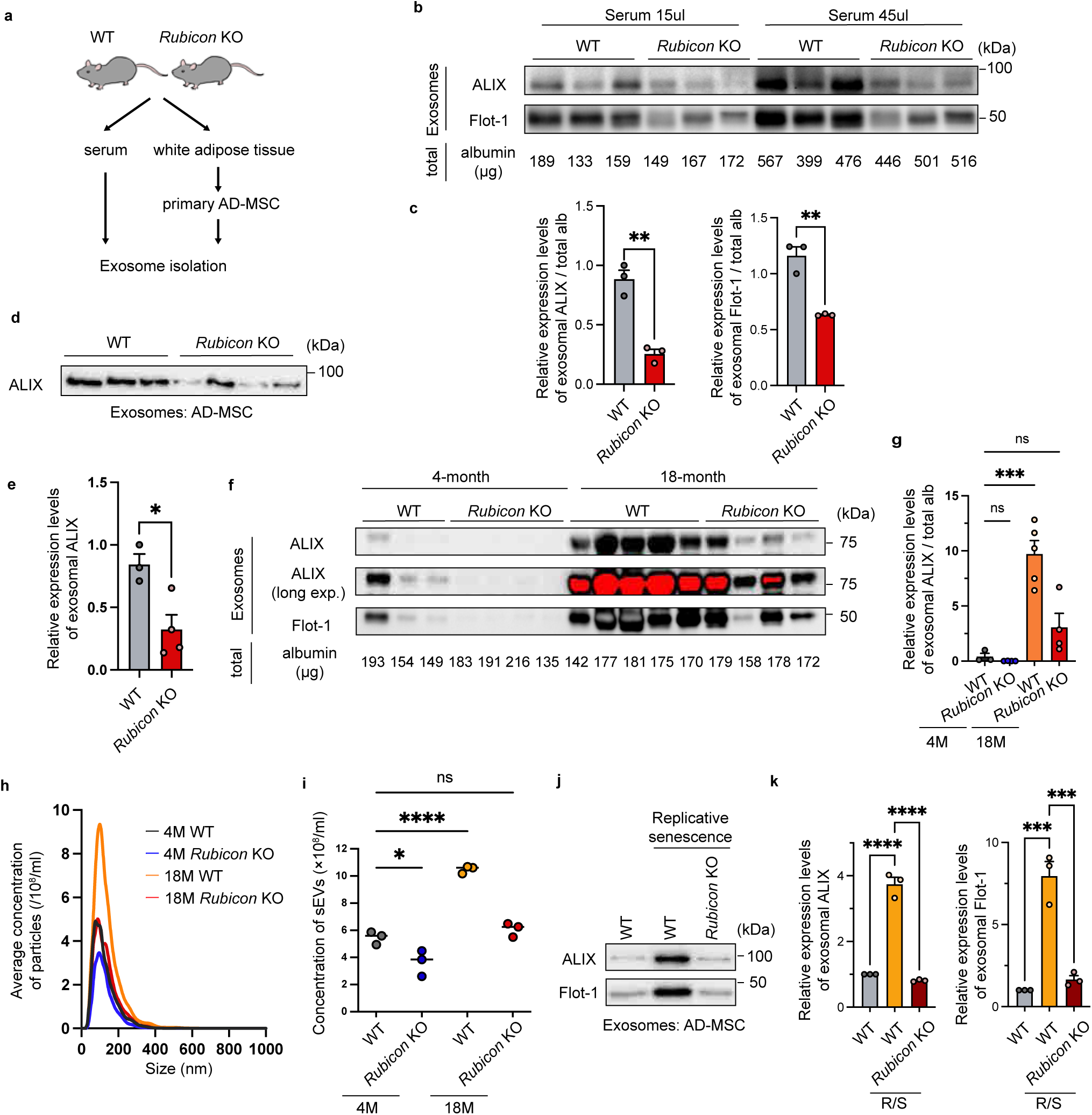
Rubicon is necessary for an age-dependent increase in serum exosomes in mice. (a) Experimental design for sampling serum or primary cells from mice. WT and *Rubicon*-KO mice were subjected to isolation of serum and white adipose tissue (WAT). Primary adipose tissue–derived mesenchymal stem cells (AD-MSCs) were obtained from the WAT using collagenases. (b) Upper: Immunoblotting of ALIX and Flot-1 in the exosome fractions purified from the 15 or 45μl serum of WT and *Rubicon-*KO mice. Bottom: Serum albumin levels of indicated mice. WT, n =3; *Rubicon*-KO, n = 3. The loading amount for each sample was set as follows: all the exosome sample collected from 15 or 45 μl of serum, respectively. (c) Quantification of the exosomal levels of ALIX and Flot-1 normalized by serum albumin levels in (b). Bars represent means ± SEM. n = 3. **p < 0.01 by a two-tailed Student’s t-test. (d) Immunoblotting of ALIX in the exosome fractions purified from the culture medium of primary AD-MSCs isolated from WT and *Rubicon*-KO mice. WT, n = 3; *Rubicon*-KO, n = 4. *p < 0.05 by a two-tailed Student’s t-test. The loading amount for each sample was set as follows: half of the exosome sample collected from 1 ml of culture medium. (e) Quantification of exosomal ALIX levels in (d). Bars represent means ± SEM. WT, n = 3; *Rubicon*-KO, n = 4. *p < 0.05 by a two-tailed Student’s t-test. (f) Upper: Immunoblotting of ALIX and Flot-1 in the exosome fractions purified from serum samples of 4-month-old or 18-month-old mice with the indicated genotypes. Bottom: Serum albumin levels of indicated mice. 4-month-old WT, n =3; 4-month-old *Rubicon*-KO, n = 4; 18-month-old WT, n = 5; 18-month-old *Rubicon* KO, n = 4. The loading amount for each sample was set as follows: all the exosome sample collected from 15 μl of serum. (g) Quantification of the exosomal ALIX levels normalized by serum albumin levels in (f). Bars represent means ± SEM. 4-month-old WT, n =3; 4-month-old *Rubicon*-KO, n = 4; 18-month-old WT, n = 5; 18-month-old *Rubicon*-KO, n = 4. ns, not significant; ***p < 0.001 by one-way ANOVA followed by Dunnett’s test. (h) NTA tracing of the EVs purified from the serum of the indicated mice. (i) Quantification of the total particle concentration of the EVs in (h). Bars represent means. n = 3. ns, not significant; *p < 0.05, ****p < 0.0001 by one-way ANOVA followed by Dunnett’s test. (j) Immunoblotting of ALIX and Flot-1 in the exosome fractions purified from the culture medium of primary AD-MSCs isolated from WT and *Rubicon*-KO mice with or without replicative senescence. n = 3. The loading amount for each sample was set as follows: half of the exosome sample collected from 1 ml of culture medium. (k) Quantification of the exosomal levels of ALIX and Flot-1 in (j). Bars represent means ± SEM. n = 3. ***p < 0.001, ****p < 0.0001 (each vs WT) by one-way ANOVA followed by Dunnett’s test.

### Rubicon is essential for the age-dependent alteration of the exosomal microRNA profiles that promote cellular senescence

Previous reports showed that exosomal miRNA packaging is mediated by ESCRT pathway.^62,63^ Therefore, we sought to determine whether Rubicon or aging influences the exosomal miRNA profile. To this end, we performed small RNA sequencing of serum exosomes isolated from the four mouse groups: young *Rubicon* wild type (WT), aged *Rubicon* WT, young *Rubicon* KO, and aged *Rubicon* KO. We identified ten miRNAs that increased with age in a manner that depends on Rubicon (Fig. 6a,b). Notably, the pathway enrichment analysis of the predicted miRNA targets showed significant enrichments in cell cycle and longevity pathways including FoxO signaling, insulin signaling and mTOR signaling (Fig. 6c and Extended Data Fig.10a). These findings suggest that Rubicon is required for the age-dependent alteration of exosomal miRNA profiles, which is involved in aging-related pathways. Because Rubicon play a pivotal role in exosome release during cellular senescence, we sought to determine whether these miRNAs could regulate cellular senescence in cultured cells. To this end, we evaluated cellular senescence markers in miRNA-overexpressing cells, and found that overexpression of miR-26a-5p or miR-486a-5p increased the levels of cellular senescence markers in IMR90 cells (Fig. 6d-g). This data suggests that the expression of miR-26a-5p and miR-486a-5p promote cellular senescence. Consistent with this, previous reports showed that miR-26a-5p and miR-486a-5p target *FOXO1*, a well-known anti-aging transcriptional factor, to suppress cell proliferation.^64,65^ Other studies showed that the miR-26a-5p targets EZH2, a histone methyltransferase that suppresses p21 expression,^66^ to promote cellular senescence.^66,67^ Overexpression of miR-26a-5p or miR-486a-5p suppressed the mRNA levels of *EZH2* and *FOXO1* in IMR90 cells (Fig. 6h). These results indicate that miR-26a-5p and miR-486a-5p, secreted into serum exosomes in a Rubicon-dependent manner during aging, suppress the expression of aging-related genes and promote cellular senescence.

**Fig. 6.**
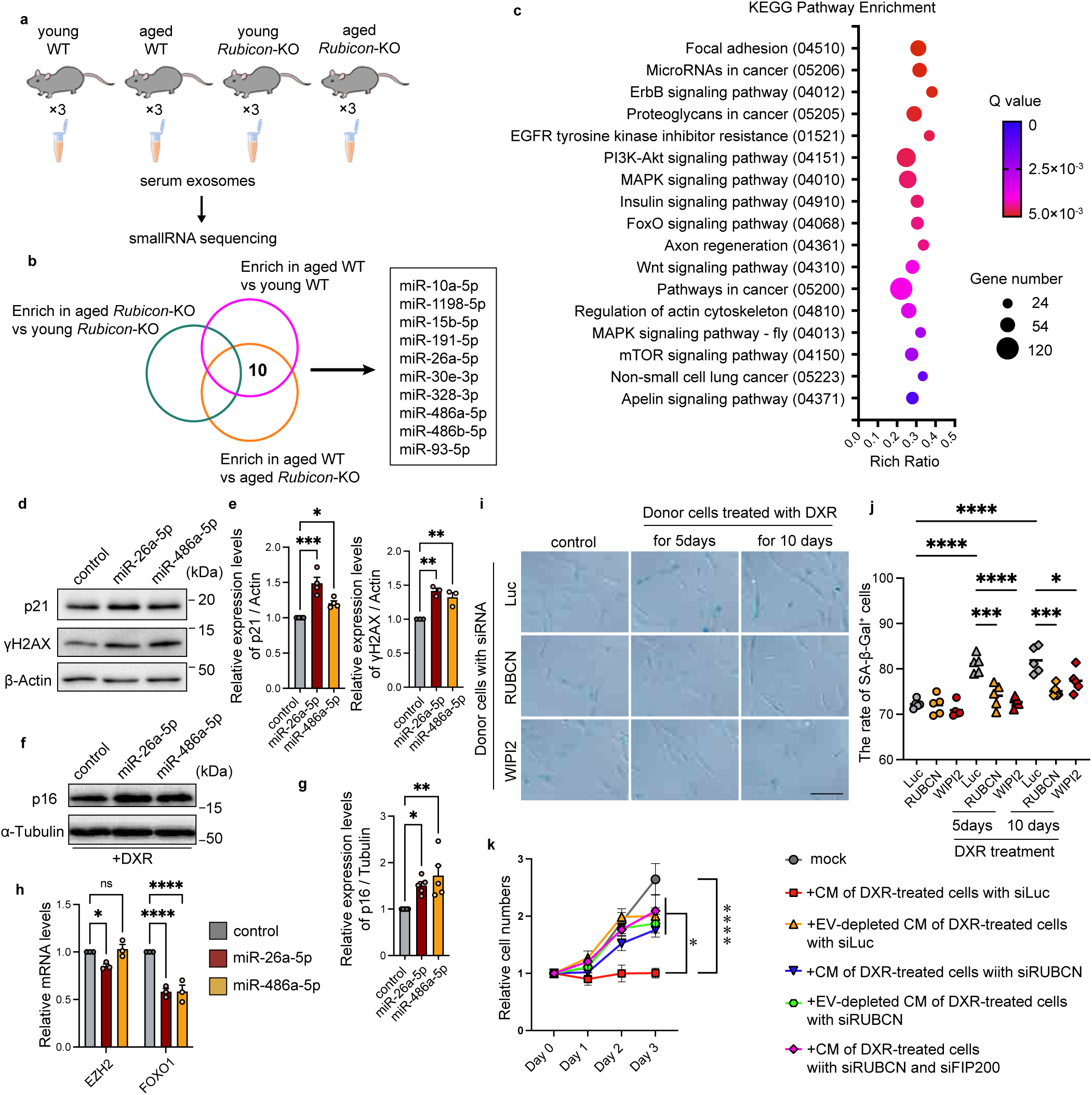
Loss of Rubicon alleviates the age-dependent alteration of the exosomal microRNA profile. (a) Illustration of the procedure for small-RNA sequencing of serum exosomes. Exosomes were isolated from the sera of 4-month-old WT (young WT), 18-month-old WT (aged WT), 4-month-old *Rubicon*-KO (young *Rubicon*-KO) and 18-month-old *Rubicon*-KO (aged *Rubicon*-KO) mice using a PS-affinity kit. n = 3 independent mice. (b) Venn diagrams showing the numbers of DEGs in serum EVs from aged WT mice relative to young WT (magenta) and aged *Rubicon*-KO (yellow) mice and aged *Rubicon*-KO relative to young *Rubicon*-KO (green). n = 3 independent mice. (c) Bubble plot exhibiting the results of Kyoto Encyclopedia of Genes and Genomes (KEGG) pathway enrichment analysis of the target genes of the 10 microRNAs (miRNAs) enriched in the serum EVs in (b). X-axis, pathway enrichment; Y-axis, pathway categories. The bubble size indicates the number of target genes. The color bar indicates the corrected q-value. (d) Immunoblotting of p21, γH2AX and β-Actin in IMR90 transfected with indicated miRNAs. P21 and β-Actin, n = 4; γH2AX, n = 3. The loading amount for each sample was set as 6 µg. (e) Quantification of levels of p21 and γH2AX normalized by β-Actin in (d). Bars represent means ± SEM. P21, n = 4; γH2AX, n = 3. *p < 0.05, **p < 0.01, ***p < 0.001 (each vs control) by one-way ANOVA followed by Dunnett’s test. (f) Immunoblotting of p16 and α-Tubulin in IMR90 transfected with indicated miRNAs and treated with doxorubicin for 15h. n = 5. The loading amount for each sample was set as 10 µg. (g) Quantification of levels of p16 normalized by α-Tubulin in (f). Bars represent means ± SEM. n = 5. *p < 0.05, **p < 0.01 (each vs control) by one-way ANOVA followed by Dunnett’s test. (h) Relative mRNA expression of EZH2 and FOXO1 in IMR90 transfected with indicated miRNAs. Bars represent means ± SEM. n = 3. ns, not significant; *p < 0.05, ****p < 0.0001 (each vs control) by one-way ANOVA followed by Dunnett’s test. (i) Representative staining images of SA-β-gal–positive recipient RPE-1 cells treated with indicated conditioned medium (CM) of donor cells. Cellular senescence in recipient RPE1 cells was induced using DXR treatment for 5 days before adding of CM. n = 5. Scale Bar, 100μm. (j) Quantification of the rate of SA-β-gal–positive cells in (i). Bars represent means. n = 5. *p < 0.05, ***p < 0.001, ****p < 0.0001 by one-way ANOVA followed by Tukey’s test. (k) Relative cell numbers of recipient IMR90 cells treated with indicated CM of donor RPE-1 cells. n = 5. *p < 0.05, ****p < 0.0001 by one-way ANOVA followed by Dunnet’s test.

### Loss of Rubicon attenuates the pro-senescent effect of extracellular vesicles secreted from senescent cells

Previous reports showed that EVs secreted by senescent cells promote cellular senescence in recipient cells^60^, thus we next asked if Rubicon-dependent EV secretion contributes to the development of cellular senescence. To this end, we employed the previously described method of adding conditioned medium (CM) of senescent donor cells to recipient cells^60^ (Extended data Fig. 10c). We found that CM of senescent donor cells increased the number of SA-β-Gal-positive cells in recipient cells (Fig. 6i,j). Importantly, this increase was abolished by knockdown of *RUBCN* or *WIPI2* in donor cells (Fig. 6i,j). Consistently, addition of CM from senescent donor cells caused growth arrest of human fibroblasts, which was rescued by EV depletion or *RUBCN* knockdown in donor cells (Fig. 6k). Of note, knockdown of an autophagy-essential gene, *FIP200*, did not affect the phenotype of *RUBCN*-knockdown donor cells (Fig. 6k). These results suggest that Rubicon mediates the development of cellular senescence through exosome secretion, but not through the autophagy pathway.

## Discussion

In the present study, we found that the Rubicon-WIPI axis positively regulates ESCRT-dependent exosome secretion. Rubicon recruits the WIPI proteins, WIPI2d and WIPI3, to endosomes, and mediates the sequential recruitment of ESCRT and associated proteins that are essential for MVB formation. Furthermore, Rubicon accumulation contributes to age-dependent changes in exosomal miRNA quantity and quality.

Recent reports suggest that non-canonical forms of autophagy, such as LAP or LANDO, require Rubicon and the WD domain of ATG16, but not WIPI2,^23,68^ which is entirely different from the Rubicon-WIPI axis in exosomal biogenesis. Importantly, our current study suggests that Rubicon regulates exosome secretion independently of Beclin-1. The potential Beclin-1-independent Rubicon complex needs to be investigated in future studies. In addition, the Rubicon-WIPI axis and the exosome secretion pathway do not require FIP200, which is indispensable for autophagosome formation; thus, both are independent of autophagosome-related pathways, including amphisome formation^27,41,42^ and autophagic lysosome reformation.^43–46^ Several studies showed that WIPI1 and its yeast homolog Atg18 mediate endosome fragmentation by localizing not only to autophagosomes but also to endosomes.^69,70^ Another report showed that ATG2, a binding partner of WIPI2 during canonical autophagy, contributes to the lysosomal damage response.^71^ However, the non-canonical function of WIPI2 was largely unknown. Our present study described the role of WIPI2 in exosome biogenesis as its non-autophagic function for the first time. However, there are still several unanswered questions regarding the mechanism involved, particularly how the initiation site of ILV formation is determined. It is conceivable that ubiquitination or endosomal lipidomes might be involved, but further research is needed. Since WIPI2 recruits ESCRT components and their associated proteins to endosomes for MVB formation, it is also plausible that WIPI2 is involved in the lysosomal recruitment of the ESCRT machinery under specific conditions, such as during the lysosomal damage response.^72,73^ This hypothesis should be tested in future studies.

Exosomes exert various physiological effects by eliminating harmful substances in donor cells^36^ and/or by transferring their cargo, such as proteins or miRNAs, to recipient cells.^3,56,59^ Our findings have shown that Rubicon- and aging-dependent exosomal miRNAs, miR-26a-5p and miR-486a-5p, suppress the expression of longevity-related genes and promote cellular senescence. Notably, recent studies showed that exosomal miRNAs secreted from young mice exhibit rejuvenate effect on recipient cells or tissues.^74^ Our small RNA sequencing also identified several exosomal miRNAs secreted from young mice, therefore, their biological effects are interesting and need to be investigated in future studies. Previous studies have shown that RBPs mediate the packaging of miRNAs into exosomes, probably via interaction with the ESCRT machinery.^63^ The Rubicon-WIPI axis controls the secretion of age-related exosomal miRNAs through the ESCRT machinery. Thus, we speculated that the Rubicon-WIPI axis mediates the interaction between this machinery and specific RBPs. Our small RNA sequencing identified exosomal miRNAs that exhibited lower expression in young or aged *Rubicon*-KO mice, including miR92a-5p and miR93-5p (Extended Data Fig. 10d). Previous reports showed that hnRNPA1 and hnRNPA2/B1 interact with miR-92a-3p or miR-93-5p for exosomal packaging.^63,75,76^ Based on these findings, we hypothesized that hnRNPA1 and hnRNPA2/B1 regulate exosomal miRNA packaging in the Rubicon-WIPI axis, and investigated whether WIPI2 was interacts with hnRNPA1 or hnRNPA2/B1. However, we were unable to detect any interaction among them (Extended Data Fig. 10e). Thus, the mechanism of exosomal miRNA packaging in the Rubicon-WIPI axis remains an open topic for future studies.

Our current results indicate that the accumulation of Rubicon is associated with age-dependent changes in exosomal contents. Moreover, our recent study showed that the accumulation of Rubicon promotes cellular senescence;^61^ therefore, the paracrine effect of exosomes on cellular senescence or longevity pathways during aging may be of particular interest. With the age-dependent decline of autophagic activity, exosomes may serve as an alternative pathway for the elimination of harmful molecules.^77^ Recent studies have reported that exosomes eliminate protein aggregates, thereby mitigating neurodegenerative diseases.^36^ Furthermore, previous reports have revealed that mutations in human Rubicon are associated with neurodegenerative diseases.^78^ It would be particularly interesting to investigate whether the Rubicon-WIPI axis is involved in the prevention of neurogenerative diseases via exosome regulation. To further investigate physiological relevance of Rubicon-dependent EVs, since the present study has focused CD63-positive exosomes, comprehensive analysis of other EVs, which are specifically detected using other EV markers, need to be performed if necessary.

## Methods

### Cell culture

hMSCs were obtained from the Lonza Inc. HEK293T, HeLa Kyoto cells and RPE-1 cells were the same as those previously used in our laboratory.^61,79,80^ *Rubicon*-KO MEFs^21^ and *RUBCN*-KO HeLa Kyoto cells^79^ were previously generated in our laboratory. *Atg16l1-KO* MEFs were previously generated by Dr. Shizuo Akira (Osaka University) in collaboration with us.^81^ *Fip200-* KO MEFs were previously described.^82^ *Atg2A-* and *Atg2B-* KO (*Atg2*-KO) MEFs, *Atg5-*KO MEFs and *Atg14*-KO MEFs were kindly gifted by Dr. Noboru Mizushima (The University of Tokyo). *Beclin-1*-KO MEFs and *Uvrag*-KO MEFs were kindly gifted by Dr. Shizuo Akira (Osaka University). *Wipi2-KO* MEFs were kindly provided by Dr. Shigeomi Shimizu (Tokyo Medical and Dental University).^83^ Plat-E cells were generously provided by Dr. Toshio Kitamura (The University of Tokyo).^84^ IMR90 cells were kindly provided by Dr. Akiko Takahashi (Japanese Foundation for Cancer Research). MEFs, HEK293T cells, HeLa Kyoto cells, Plat-E cells, IMR90 cells and RPE-1 cells were cultured in Dulbecco’s Modified Eagle’s medium (Sigma-Aldrich, D6046) supplemented with 10% fetal bovine serum (Gibco, 10270106) and 1% penicillin– streptomycin (Sigma-Aldrich, P4333). The MSCs were cultured in Mesenchymal Stem Cell Growth Medium 2 (PromoCell, C-28009) supplemented with 1% penicillin–streptomycin. RPE-1 cells were incubated with 150 ng/mL doxorubicin (FUJIFILM Wako Chemicals, 040-21521) for 5 or 10 days to induce DNA damage-induced cellular senescence. In replicative senescence, late passage mouse MSCs (> 70 population doublings, PDL) were used as senescent cells and early passage mouse MSCs (< 30 PDL) were used as controls.

### Mouse studies

Whole-body *Rubicon*-KO mice on a C57BL/6 background were generated as described previously.^22^ Mice were housed in box cages, maintained on a 12-hour light / 12-hour dark cycle, and fed a normal-fat chow diet (Oriental Yeast, MF). The ambient temperature and humidity were 23°C ± 1.5°C and 45% ± 15%, respectively. All animal experiments were approved by the Animal Research Committee of Osaka University and complied with the Japanese Animal Protection and Management Law (No. 25). We complied with all relevant ethical regulations.

### Reagents and antibodies

The following antibodies were used for western blotting at the indicated dilutions: anti-CD63 (MBL, D263-3, 1:5000; Abcam, ab134045, 1:2000), anti–Flotilin-1 (BD Biosciences, 610821, 1:10000), anti-ALIX (BioLegend, 634502, 1:1000), anti-Calnexin (Sigma-Aldrich, HPA009433, 1:1000), anti-Rubicon (Cell Signaling Technology, 8465, 1:2000), anti–α-Tubulin (PM054, MBL, 1:20000), anti-WIPI2 (Sigma-Aldrich, SAB4200400, 1:2000), anti-HRS (GeneTex, GTX101718, 1:1000), anti-VPS4 (Sigma-Aldrich, SAB4200025, 1:1000), anti-GFP (Cell Signaling Technology, 2555, 1:1000), anti-p21 (Abcam, ab109199, 1:1000), anti-γH2AX (Abcam, ab2893, 1:1000), anti-p16 INK4a (IBL, 11104, l g/ml), anti-LC3 (MBL, PM036, 1:2000), anti-Atg12 (Cell Signaling Technology, 2011S, 1:1000), anti-FIP200 (Proteintech, 17250-1-AP, 1:1000), anti-Atg2A (MBL, PD041, 1:1000), anti-Atg2B (Sigma-Aldrich, HPA019665, 1:1000), anti-Atg14 (Cell Signaling Technology, 96752S, 1:1000), anti–Beclin-1 (Cell Signaling Technology, 3738S, 1:1000), anti-UVRAG (MBL, M160-3, 1:2000), anti-ATG16L1 (Cell Signaling Technology, 8089, 1:1000), anti-PIK3C3 (Cell Signaling Technology, 3811, 1:1000), anti-WIPI1 (Sigma-Aldrich, W4769, 1:1000), anti-WIPI3 (Thermo Fisher Scientific, PA5-50864, 1:1000), anti-WIPI4 (Proteintech, 19194-1-AP, 1:1000), anti–β-Actin (MBL, M177-3. 1:2000), anti-FLAG (Sigma-Aldrich, F1804, 1:5000), anti-HA (Roche, 12158167001, 1:3000), anti-CHMP4B (Proteintech, 13683-1-AP, 1:1000), anti-TSG101 (BD Biosciences, 612696, 1:1000), anti-EGFR (Fitzgerald, 20-ES04, 1:1000), horseradish peroxidase (HRP)-conjugated goat anti–rabbit IgG (Jackson ImmunoResearch, 111–035-003, 1:5000), HRP-conjugated goat anti–rat IgG (Jackson ImmunoResearch, 112–035-003, 1:5000), and HRP-conjugated goat anti–mouse IgG (Jackson ImmunoResearch, 115–035-003, 1:5000). The following antibodies were used for immunofluorescence: anti-CD63 (MBL, D263-3, 1:500; BD Biosciences, 556019, 1:1000), anti-LAMP1 (BD Biosciences, 553792, 1:1000), anti-WIPI2 (Sigma-Aldrich, SAB4200400, 1:500), anti-WIPI1 (Sigma-Aldrich, W4769, 1:200), anti-WIPI3 (Thermo Fisher Scientific, PA5-50864, 1:200), anti-ALIX (BioLegend, 634502, 1:200), goat anti–mouse IgG H&L Alexa Fluor 488 pre-adsorbed (Abcam, ab150117, 1:2000), goat anti–rat IgG H&L Alexa Fluor 568 pre-adsorbed (Abcam, ab175710, 1:2000), goat anti–rabbit IgG H&L Alexa Fluor 568 pre-adsorbed (Abcam, ab175695, 1:2000), goat anti–mouse IgG (H+L) cross-adsorbed secondary antibody, Alexa Fluor 568 (Invitrogen, A11004, 1:2000), and goat anti–rat IgG H&L Alexa Fluor 647 pre-adsorbed (Abcam, ab150167, 1:2000). Apilimod and bafilomycin A_1_ were purchased from MedChemExpress (HY-14644) and Wako (023-11641), respectively.

### Plasmid construction

An empty pcDNA3.1 vector was purchased from Invitrogen (V79020). A pEGFP-hCD63 vector was purchased from Addgene (62964). Empty pMRX-IRES-bsr and pMRX-IRES-puro vectors were kindly gifted by Dr. Shoji Yamaoka (Tokyo Medical and Dental University). pMRX-puro-ATG16L1 vectors, including the full-length (FL;1–607) and ΔWD repeat domain (ΔWDR; 1– 249),^82^ pcDNA3.1-3×FLAG-hRubicon,^85,86^ and pMRX-bsr-GFP-mRubicon^21^ were previously generated. The mutant 3×FLAG-tagged hRubicon vectors were generated using an in-Fusion reaction (TaKaRa Bio). The mCherry-tagged mRubicon was subcloned into pMRX-IRES-bsr. The mStrawberry-tagged WIPI2d, GFP-tagged WIPI2a and WIPI2d, mCherry-tagged 2×FYVE and GFP-tagged hCD63 were subcloned into pMRX-IRES-puro. The HA-tagged WIPI2a, WIPI2b, WIPI2c, WIPI2d, hnRNPA1, hnRNPA2 and hnRNPB1 were subcloned into pcDNA3.1. The GFP-tagged VPS4 was subcloned into pcDNA3.1 and modified to pcDNA3.1-GFP-VPS4E228Q using in-Fusion reaction. The GFP-tagged Rab5 was subcloned into pcDNA3.1 and modified to pcDNA3.1-GFP-Rab5Q79L using an in-Fusion reaction. pCX-bG-mmu-miR-26a and pCX-bG-mmu-miR-486a were generated as described previously^87^. Briefly, the miR-26a and miR-486a gene loci were amplified from mouse genomic DNA and ligated into a pCX-bG vector, respectively. Recombinant retroviruses were prepared as previously described.^88^ Stable transformants were selected with 10 μg/mL blasticidin or 1.5 μg/mL puromycin. Plasmid transfections were performed using the Lipofectamine^TM^ 2000 Transfection Reagent (Thermo Fisher Scientific, 11668500).

### RNA interference

The following siRNA duplex oligomers were generated by Sigma-Aldrich as the indicated sense strands: 5′-UCGAAGUAUUCCGCGUACGdTdT-3′ for *Luciferase*; 5′-CGCCUGUUCUACGAGAAGAdTdT-3′ for *ULK1*; 5′-UAUCACAUGAGUGGU CUGAdTdT-3′ for *ULK2*; 5′-UAGGAACUCCAACAACUGGGAAUGC-3′ for *ATG2A*; 5′-UAAAGAUGCCAGACAAGAGAGACCU-3′ for *ATG2B*; 5′-CAUUGAGACUGUUGCAGAAdTdT-3′ for *ATG3*; 5′-UAAAGCAGUACAUACAGGCdTdT-3′ for *ATG5*; 5′-UUGUUGGACGUCUUAGACCdTdT-3′ for *BECN1*; 5′-GGAGUCACAGCUCUUCCUUdTdT-3′ for *ATG7*; 5′-CAAGAAGCAAUUCAUGUACdTdT-3′ for *ATG9*; 5′-GCGUAAUAGUGUCCCAUGGdTdT-3′ for *ATG10*; 5′-GCAGUAGAGCGAACACGAAdTdT-3′ for *ATG12*; 5′-GAGUUUGGAUAUACCCUUUdTdT-3′ for *ATG13*; 5′-GCAAGAUGAGGAUUGAACAdTdT-3′ for *ATG14*; 5′-AGCTTCAAATGATTTTGCAdTdT-3′ for *ATG16L1;* 5′-GAUCUUAUGUAGUCGUCCAdTdT-3′ for *FIP200*; 5′-GUCAGAAUGAGGAUGACAUdTdT-3′ for *RUBCN*; 5’-AAACUCAACACUGGCUAAUUAdTdT-3’ for *PIK3C3* (#2); 5’-GCACAAACCCAGGCUAUUUdTdT-3’ for *PIK3C2α*; 5’-GCCGGAAGCUUCUGGGUUUdTdT-3’ for *PIK3C2β*; 5’-GAUCGAAAGCCAAAUAGUUdTdT-3’ for *INPP4a*; 5’-CGAUGAAAUUGGAAUGUUAdTdT-3’ for*INPP4b*; 5’-CUGUGGUUUGCAUGUCGGAdTdT-3’ for *VPS4A*; 5’-CCAAAGAAGCACUGAAAGAdTdT-3’ for *VPS4B*; 5’-GAAGGAUGCUUUCGAUAAAdTdT-3’ for *ALIX*; 5’-CCUCCAGUCUUCUCUCGUCdTdT-3’ for *TSG101* and 5’-CGACAAGAACCCACACGU-dTdT-3’ for *HRS*. The *ATG4A* (SASI_Hs01_00150665), *ATG4B* (SASI_Hs01_00126888), *ATG4C* (SASI_Hs01_00075254), *ATG4D* (SASI_Hs01_00023430), and *CHMP4B* (SASI_Hs01_00129003) siRNAs were purchased from Sigma-Aldrich. The ON-TARGETplus siRNA SMART pools for Non-targeting Control (D-001810-10), *WIPI2* (L-020521-01), and other candidate genes in the RNAi screen (Fig. 2a and S5a) were purchased from Dharmacon. The Silencer™ Negative Control No. 1 (AM4611), *WIPI1* (s30081), *WIPI2* (#2) (s25100), *WIPI3* (s32119), and *WIPI4* (s533528) siRNAs were purchased from Thermo Fisher Scientific. The hMSCs or HEK293T cells were transfected with siRNAs using Lipofectamine^TM^ RNAiMAX Transfection Reagent (ThermoFisher Scientific, 13778150) and used for the subsequent experiment 48 hours after transfection.

### RNA sequencing

Library preparation and sequencing were conducted by BGI. Small RNA with a size of 18–30 nt was isolated by gel electrophoresis. Libraries were prepared with the MGIEasy Small RNA Library Prep Kit (940-000196-00, MGI), and quality and quantity were assessed at all steps by capillary electrophoresis (Agilent Bioanalyzer). Libraries were sequenced on a DNBSEQ-G400 with cPAS chemistry (DNBSEQ-G400RS High-throughput Sequencing Set, FCL SE50, 1000016941, MGI).

### Western blotting

Cells were lysed in 20 mM Tris-HCL pH 7.4, 150 mM NaCl, 1 mM EDTA, 1% Triton-X100 (Nacalai Tesque, 35501-15), 1% sodium deoxycholate (Sigma-Aldrich, D6750), 0.1% SDS, 1 mM PMSF, and protease inhibitor mixture (Roche, 11873580001) on ice. After centrifugation at 20,380 × g for 15 minutes, the protein concentrations in the supernatant were measured using a BCA assay (Nacalai Tesque, 06385-00). Lysates were dissolved with non-reducing buffer (30% glycerol, 50 mM Tris-HCl, 10% SDS, 10 mM EDTA, 0.1% bromophenol blue, pH 6.8) for detecting mouse CD63, or with reducing sample buffer (30% glycerol, 50 mM Tris-HCl, 10% SDS, 250 mM dithiothreitol, 10 mM EDTA, 0.1% bromophenol blue, pH 6.8) for detecting other proteins. Protein lysates were separated by SDS-PAGE gels and transferred to PVDF membranes (Merck, IPVH00010). Membranes were stained with Ponceau-S, blocked with 3% skim milk TBS-T (tris-buffered saline and 0.1% Tween 20) for 30 minutes, incubated overnight at 4°C with a primary antibody in 3% skim milk TBS-T, and then incubated with an HRP-conjugated secondary antibody in 3% skim milk TBS-T for 1 hour. Blots were developed with an Immobilon Forte Western HRP substrate (Merck, WBLUF0500) or an ImmunoStar LD (Fujifilm Wako, 290-69904), and signals were detected using a ChemiDoc^TM^ Touch Imaging System (Bio-Rad).

### Exosome isolation

Exosome isolation from the cell culture medium was performed as previously described.^59^ Briefly, the MEFs or MSCs were cultured with medium containing 10% exosome-depleted FBS or Serum-Free Medium STK2^®^ for Mesenchymal Stem Cell (Twocells, 37415-08), respectively, for the indicated durations. The culture medium was centrifuged, and the resultant supernatants were ultracentrifuged at 110,000 × g using a Fixed-Angle Rotor TLA55 (Beckman Coulter, 366725) followed by washing of the exosome pellets with phosphate-buffered saline (PBS). The exosome pellets were directly solubilized in the sample buffer for immunoblotting. Exosome isolation from mouse serum was performed using an ExoQuick precipitation reagent (System Biosciences, EXOQ5A-1) and a Fixed-Angle Rotor TLA120.1 (Beckman Coulter, 362224) as previously described.^59^ Serum albumin levels in total fraction were measured using QuantiChrom^TM^ BCG Albumin Assay Kit (BioAssay Systems, DIAG-250). A MagCapture™ Exosome Isolation Kit PS Ver.2 (Fujifilm Wako, 290-84103) was used to isolate exosomes for NTA assays. For comparative analysis, exosomes were collected from equal volumes of culture medium and conditioned by equal numbers of cells. Immunoblotting showed that the isolated exosome fraction was positive for multiple exosome markers and did not contain any organelle markers. The density and size distribution of exosomes were analyzed using a NanoSight LM10-HS apparatus and NTA2.3 software (Malvern Panalytical).

### Immunofluorescence staining

After fixation with 4% paraformaldehyde (Nacalai Tesque, 09154-85) for 20 minutes at room temperature, cells were treated with 50 μg/mL digitonin in PBS for 10 minutes at room temperature. The cells were incubated with 0.1% gelatin in PBS for 30 minutes at room temperature, and then reacted with primary antibodies for 60 minutes at room temperature. After incubation with a fluorescence-labeled secondary antibody for 40 minutes at room temperature, cells were covered with a VECTASHIELD® Antifade Mounting Medium with DAPI (VECTOR laboratories, H-1200), and imaged with an FV3000 Confocal Laser Scanning Microscope (Olympus) operated using FV31S-SW software (version 2.3.1.163).

### Gel filtration

MEFs and hMSC lysate were subjected to gel filtration as previously described^37^ with some modifications. Briefly, cells cultured on 15 cm dishes were harvested, washed with PBS, and then lysed for 15 min at 4°C in a solution containing 20 mM HEPES-KOH (pH 7.5), 150 mM NaCl, 1 mM MgCl_2_, 0.2% Triton X-100, 1 mM PMSF, 2 μg/ml pepstatin and 100 µg/mL protease inhibitor cocktail (Roche). The lysate was clarified by ultracentrifugation using a TLA-55 rotor (Beckman Coulter) at 100,000 × g for 10 min at 4°C and also by centrifugal filtration using Centricut Ultramini MF (0.2 µm) units (Kurabo). Subsequently, 250-400 µL of the clarified sample was combined with 10-30 µL of a molecular weight marker solution prepared from Gel Filtration Standard (Bio-Rad) according to the manufacturer’s instructions, loaded onto a Superdex 200 Increase 10/300 column (Cytiva) pre-equilibrated with 20 mM HEPES-KOH (pH 7.5), 150 mM NaCl, and 1 mM MgCl_2_ using the ÄKTA pure 25 M1 chromatography system (Cytiva), and then eluted with the same buffer at a flow rate of 0.4 mL/min while collecting every 0.2 mL of eluate. A selected range of these fractions was finally analyzed by Western blotting.

### Immunoprecipitation and MS using MEFs

MEFs stably expressing the indicated plasmids were grown in a 15-cm dish and treated with or without Apilimod for 1 hour, and then cross-linked with 0.1% formaldehyde (Wako, 063-04815) for 10 minutes at room temperature. After cross-linking was quenched with 100 mM glycine for 4 minutes at room temperature, cells were washed with HEPES saline containing 20 mM HEPES-NaOH buffer (pH 7.5) and 137 mM NaCl, and lysed in HEPES-RIPA buffer containing 20 mM HEPES-NaOH buffer (pH 7.5), 150 mM NaCl, 1 mM EGTA, 1 mM MgCl_2_, 0.25% (w/v) sodium deoxycholate, 0.05% SDS, 1% (v/v) NP-40, 0.2% (v/v) Benzonase Nuclease (Millipore, 70746), PhosSTOP (Roche, 4906837001), and Protease Inhibitor Cocktail (Merck, 11873580001) on ice. After sonication and centrifugation at 20,380 × g at 4°C for 15 minutes, the supernatants were incubated with GFP-Trap magnetic agarose beads (ChromoTek, gtma-20) at 4°C for 2 hours. The beads were washed four times with HEPES-RIPA buffer and then twice with 50 mM ammonium bicarbonate. Proteins on the beads were digested by adding 200 ng of a trypsin/Lys-C mix (Promega) at 37°C overnight. The resultant digests were reduced, alkylated, acidified, and desalted using a GL-Tip SDB. The eluates were evaporated and dissolved in 0.1% trifluoroacetic acid and 3% acetonitrile (ACN). LS-MS/MS analysis of the resultant peptides was performed on an EASY-nLC 1200 UHPLC connected to an Orbitrap Fusion mass spectrometer through a nanoelectrospray ion source (Thermo Fisher Scientific). The peptides were separated on a C18 reversed-phase column (75 μm × 150 mm; Nikkyo Technos) with a linear 4–32% ACN gradient for 0–100 minutes, followed by an increase to 80% ACN for 10 minutes and a final hold at 80% ACN for 10 minutes. The mass spectrometer was operated in data-dependent acquisition mode with a maximum duty cycle of 3 seconds. MS1 spectra were measured with a resolution of 120,000, an automatic gain control target of 4e5, and a mass range of 375–1,500 m/z. HCD MS/MS spectra were acquired in the linear ion trap with an automatic gain control target of 1e4, an isolation window of 1.6 m/z, a maximum injection time of 35 milliseconds, and a normalized collision energy of 30. Dynamic exclusion was set to 20 seconds. Raw data were directly analyzed against the SwissProt database restricted to *Mus musculus* using Proteome Discoverer 2.5 (Thermo Fisher Scientific) with the Sequest HT search engine. The search parameters were as follows: (a) trypsin as an enzyme with up to two missed cleavages; (b) precursor mass tolerance of 10 ppm; (c) fragment mass tolerance of 0.6 Da; (d) carbamidomethylation of cysteine as a fixed modification; and (e) acetylation of the protein N-terminus and oxidation of methionine as variable modifications. Peptides were filtered at a false discovery rate of 1% using the Percolator node. Label-free quantification was performed on the basis of the intensities of precursor ions using the Precursor Ions Quantifier node. Normalization was performed such that the total sum of abundance values for each sample over all peptides was the same.

### Immunoprecipitation and MS using HeLa Kyoto cells

HeLa Kyoto cells transiently expressing the indicated plasmids were grown in a 10-cm dish with or without starvation treatment using Earle’s Balanced Salt Solution (Sigma-Aldrich, E2888). Proteins bound to the beads were reduced with 10 mM TCEP and alkylated with 20 mM IAA. Proteins were mixed with sample/methanol/chloroform/water (1:4:1:3, v/v/v/v) and spun at 14,000 × g for 5 minutes at 30°C to separate aqueous and organic layers. After the aqueous layer was removed, the organic layer was added four times the volume of methanol and spin at 21,880 × g at 30°C for 15 minutes. Proteins removed from the supernatant were dissolved in 50 µL of 9 M urea and then 8 M urea buffer diluted with 50 mM NH_4_HCO_3_ until the concentration was <1 M urea. Proteins were digested by adding 200 ng of Trypsin Gold, Mass Spectrometry Grade (Promega) at 30°C for 17 hours. The digests were acidified with 1% trifluoroacetic acid, and desalted using a SPE C-TIP following the procedure manual (Nikkyo Technos). The eluates were evaporated and dissolved in 15 µL of 1% trifluoroacetic acid and 2% acetonitrile. The 12 µL of samples were analyzed on an UltiMate 3000 UHPLC connected to a Q-Exactive Plus mass spectrometer (Thermo Fisher Scientific) through a DreamSpray ion source (AMR INCORPORATED) in data-dependent acquisition mode with the top-10 MS/MS method. MS1 spectra were measured at a resolution of 70,000, an automatic gain control target of 1e6, and a mass range from 300 to 1,500 m/z. HCD MS/MS spectra were triggered at a resolution of 17,500, an automatic gain control target of 2e5, an isolation window of 1.6 m/z, a maximum injection time of 100 milliseconds, and a normalized collision energy of 27. Dynamic exclusion was set to 20 seconds. Protein identification from raw data was performed against the SwissProt database (2017 Oct.) restricted to *Homo sapiens* using Sequest HT and the Mascot search engine on Proteome Discoverer 2.2 (Thermo Fisher Scientific). The search parameters were as follows: (a) trypsin as an enzyme with up to three missed cleavages; (b) precursor mass tolerance of 10 ppm; (c) fragment mass tolerance of 0.02 Da; (d) carbamidomethylation of cysteine as a fixed modification; and (e) acetylation of the protein N-terminus, oxidation of methionine, deamidation of asparagine and glutamine, and phosphorylation of serine, threonine, and tyrosine as variable modifications.

Peptides and proteins were filtered at a false discovery rate of 1% using the Percolator node. Label-free quantification was performed based on intensities of precursor ions using the Precursor Ions Quantifier node. Normalization was performed as the total peptide amount for each sample over all peptides was the same.

### Co-immunoprecipitation

HEK293T or HeLa Kyoto cells expressing the indicated tagged proteins were washed twice with ice-cold PBS and lysed in M-PER mammalian protein extraction reagent (Thermo Fisher Scientific, 78501) containing protease and phosphatase inhibitors. The soluble fractions from the cell lysates were obtained after centrifugation at 15,000 × g for 10 minutes. The soluble lysates were incubated with anti-FLAG M2 magnetic beads (Sigma-Aldrich, M8823) or Pierce™ Anti-HA Magnetic Beads (Thermo Fisher Scientific, 88836) with constant rotation at 4°C for 2 hours. After the incubation, the beads were washed four times with ice-cold TBS-T. For the GFP-trap assay, the cultured cells were cross-linked, lysed, and sonicated using the same method as detailed in the section titled *Immunoprecipitation and MS using MEFs*. After sonication, the supernatants were incubated with GFP-trap agarose (ChromoTek, gta-20) overnight at 4°C and washed six times with HEPES-RIPA buffer. The samples were eluted with SDS buffer and incubated at 90°C for 5 minutes, and then analyzed by immunoblotting using the indicated antibodies.

### Electron microscopy

Cells were fixed with 2.5% glutaraldehyde (Fujifilm Wako, 073-00536) in PBS and then in 2% OsO_4_ solution. They were embedded in Quetol812 (Nisshin EM), and then 80-nm ultrathin sections were obtained using an Ultracut E ultramicrotome (Reichert-Jung). These sections were stained with a solution of uranyl acetate and lead, and observed using a transmission electron microscope (Hitachi, HT-7800).

### Immunoelectron microscopy

Cells were cultured on a polystyrene cover slip, Cell Desk (Sumitomo Bakelite Co., Ltd., Japan), fixed for 1 h with 4% formaldehyde in 0.1M phosphate buffer (pH7.4) and washed for 5 min three times in 0.1M phosphate buffer (pH7.4) containing 4% sucrose. Cells were permeabilized with 0.1 M phosphate buffer (pH 7.4) containing 0.25 % saponin for 30 min and blocked with 10% BSA, 10% Normal Goat serum and 0.1% cold water fish skin gelatin in the same buffer. Cells were stained with 1:200 diluted primary antibody, anti-GFP antibodies (Nacalai Tesque, GF090R) over night at 4℃, washed for 10 min 6 times in 0.1M phosphate buffer (pH7.4) containing 0.005% saponin and followed by staining with 1:400 diluted anti Rat IgG antibodies conjugating 1.4 nm gold particle, Nanogold -Fab’ fragment of goat anti rabbit IgG (Nanoprobes Inc., USA) for 2 h at room temperature, washed for 10 min 5 times in the same buffer containing 0.005% saponin and for another 10 min without saponin. Cells were fixed with 1% (w/v) glutaraldehyde in 0.1M PB for 10 min at room temperature. Cells were washed in PBS containing 50 mM glycine, followed by washing in PBS containing 1% BSA in water. Gold labeling was intensified with GoldEnhance EM kit (Nanoprobes) for 5 min. The gold intensification solution was removed, the sections were soaked in 1% sodium thiosulfate solution for a few seconds, and washed in water. The cells were post-fixed in 1% OsO_4_ and 1.5% potassium ferrocyanide in 0.1 M phosphate buffer (pH 7.4) for 1 h. cells were dehydrated in a graded series of ethanol and embedded in epoxy resin. 80 nm ultrathin sections were stained with 8% uranyl acetate and lead staining solution. The samples were examined using a JEM-1400 plus electron microscope (JEOL, Tokyo, Japan) at 80 kV with a CCD Veleta 2K × 2K camera (Olympus).

### Isolation of primary AD-MSCs

eWAT was minced in 20 mL of Krebs-Ringer-bicarbonate-HEPES (KRBH) buffer (120 mM NaCl, 4 mM KH_2_PO_4_, 1 mM MgSO_4_·7H_2_O, 1 mM CaCl_2_, 10 mM NaHCO_3_, 30 mM HEPES, 20 µM adenosine, and 4% BSA). After centrifugation, 500 μL of 3 mg/mL Collagenase (Sigma-Aldrich, C6885) was added to the supernatant and the mixture was incubated at 37°C for 45 minutes under constant shaking. The cell suspension was filtered through a 100-μm cell strainer and then centrifuged to separate the pellet from the floating mature adipocyte fraction. The cells were washed three times, then resuspended and cultured with Mesenchymal Stem Cell Growth Medium 2 (PromoCell, C-28009).

### SA-β-Gal staining

SA-β-Gal staining was performed using the Senescence Cells Histochemical Staining Kit (Sigma, CS0030) as described previously^61^. Briefly, senescence-induced cells cultured on collagen-coated coverslips were fixed with Fixation buffer for 7 min at room temperature. After washing with PBS three times, cells were stained with Staining mixture at 37°C for 19 h without CO_2_ and then observed using a microscope (BX53, Olympus) operated using a cellSens platform (version 1.16).

### RNA analysis

Total RNA was extracted from IMR90 cells with ISOGEN II (Nippon gene). To quantify the expression levels of mRNAs, 1 μg of total RNA was reverse-transcribed with iScript^TM^ cDNA Synthesis Kit (BIO-RAD, 1708891). Quantitative PCR were performed using Power SYBR^®^ Green PCR Master Mix (Thermo Fisher Scientific, 4367659). Targets were measured using QuantiStudio^TM^ Real-Time PCR Software version 1.3. (Thermo Fisher Scientific) with the following primers: *EZH2* forward 5′-AGTTTGCTGCTGCTCTCAC-3′, reverse, 5′-GTTCTCTCCCCCCGTTTC-3′; *FOXO1* forward 5′-GCTGCATCCATGGACAACAACA-3′, reverse, 5′-CGAGGGCGAAATGTACTCCAGTT-3′; *GAPDH* forward 5′-TGTGTCCGTCGTGGATCTGA-3′, reverse, 5′-CCTGCTTCACCACCTTCTTGAT-3′; *Rnu6* forward, 5′-C GCTTCGGCAGCACATATAC-3′, reverse, 5′-TGCGTGT CATCCTTGCGCAG-3′. *GAPDH* was used as an internal control. To quantify the miRNAs, ten ng of total RNA was reverse transcribed with the TaqMan Advanced miRNA cDNA Synthesis Kit (A28007). The expression levels of mature miRNAs were quantified with real-time quantitative PCR using the TaqMan Fast Advanced Master Mix (4444557) and TaqMan MicroRNA Assays for each miRNA. *Rnu6* was used as an internal control against which to normalize the miRNA levels.

### Processing of RNA-sequencing data

Raw reads were trimmed and filtered using SOAPnuke and then aligned to reference using Bowtie2 (v2.2.9). The small RNA expression level was calculated by counting absolute numbers of molecules using unique molecular identifiers.^89^ Differential gene expression analysis was conducted using DEGseq (ver 1.30.0) with a Q-value < 0.001. The gene ontology and KEGG enrichment analyses were performed using the Dr. Tom system (BGI) with a Q-value < 5.0×10^-3^ or −log_10_ (Q value) > 2.5, respectively. Target prediction was conducted with multiple software platforms, including miRanda (v3.3a),^90^ RNAhybrid (v0.1),^91,92^ and TargetScan (v7.1).^93^

### Image analysis

Protein levels in the immunoblots and the filing rate or the dot number of CD63 in GFP-Rubicon– or GFP-Rab5Q79L–decorated endosomes were calculated using Fiji (ImageJ version 2.9.0/1.53t).

### Statistical analysis and reproducibility

All analyses were performed using GraphPad Prism 9 software (GraphPad). Statistical analyses were performed as described in the text using a two-tailed Student’s *t*-test, or one- or two-way ANOVA followed by Dunnett’s or Tukey’s multiple comparisons test. Data were collected from biologically independent experiments or samples and were denoted as means ± SEM.

## Acknowledgements

We thank Dr. Shigeomi Shimizu (Tokyo Medical and Dental University) for providing us with *Wipi2-*KO MEFs and Dr. Atsushi Nakagawa (Osaka University) for materials and technical assistance in gel filtration experiments. We thank Dr. Giovanni Bravin (Osaka University) for assistance in gel filtration experiments. We also thank the Center for Medical Research and Education and the CoMIT Omics Center for their technical support and access to the experimental equipment. T. Yoshimori is supported by JST CREST (grant no. JPMJCR17H6), JSPS KAKENHI (22H04982), and AMED (grant no. JP21gm5010001). S.N. is supported by AMED-PRIME (20gm6110003h0004). T.K. is supported by JSPS KAKENHI (20K05839). K.Y. is supported by JSPS KAKENHI (22J12444).

## Author contributions

K.Y., A.K. and T. Yoshimori planned the study and designed experiments. M.H., K.T., R.K., and T.T. conducted interactome experiments using HeLa Kyoto cells. S.K., I.S., and R.H. supervised exosome-related experiments. T. Yamamuro conducted mouse experiments. K.N. and H.K. conducted interactome experiments using MEFs. S.N. assisted with experiments related to *ATG* KO cells. H.O. performed immunoelectron microscopy. T.K. conducted gel filtration experiments. S.O., Y.K. and T.A. generated plasmids of microRNA experiments and assisted with them. R.E. and Y.O. assisted RNA sequencing. M.I. and N.B. assisted with experiments involving cultured cells. Y.T. and A.T. assisted with experiments related to aging. S.N. and S.Y. provided general assistance with this manuscript. K.Y., T. Yamamuro, S.K., and T. Yoshimori wrote the manuscript.

## Data availability

The MS proteomics data have been deposited in the ProteomeXchange Consortium via the jPOST partner repository with the dataset identifiers PXD043695, PXD043701 and PXD051908, respectively. RNA sequencing data are deposited in the National Center for Biotechnology Information Gene Expression Omnibus (https://www.ncbi.nlm.nih.gov/geo/), with accession number GSE262794.

## Competing interest

T. Yoshimori is the founder of AutoPhagyGO.

## Correspondence and materials availability

Requests for materials of this study should be addressed to Tamotsu Yoshimori.

---

**Extended Data Fig. 1.**
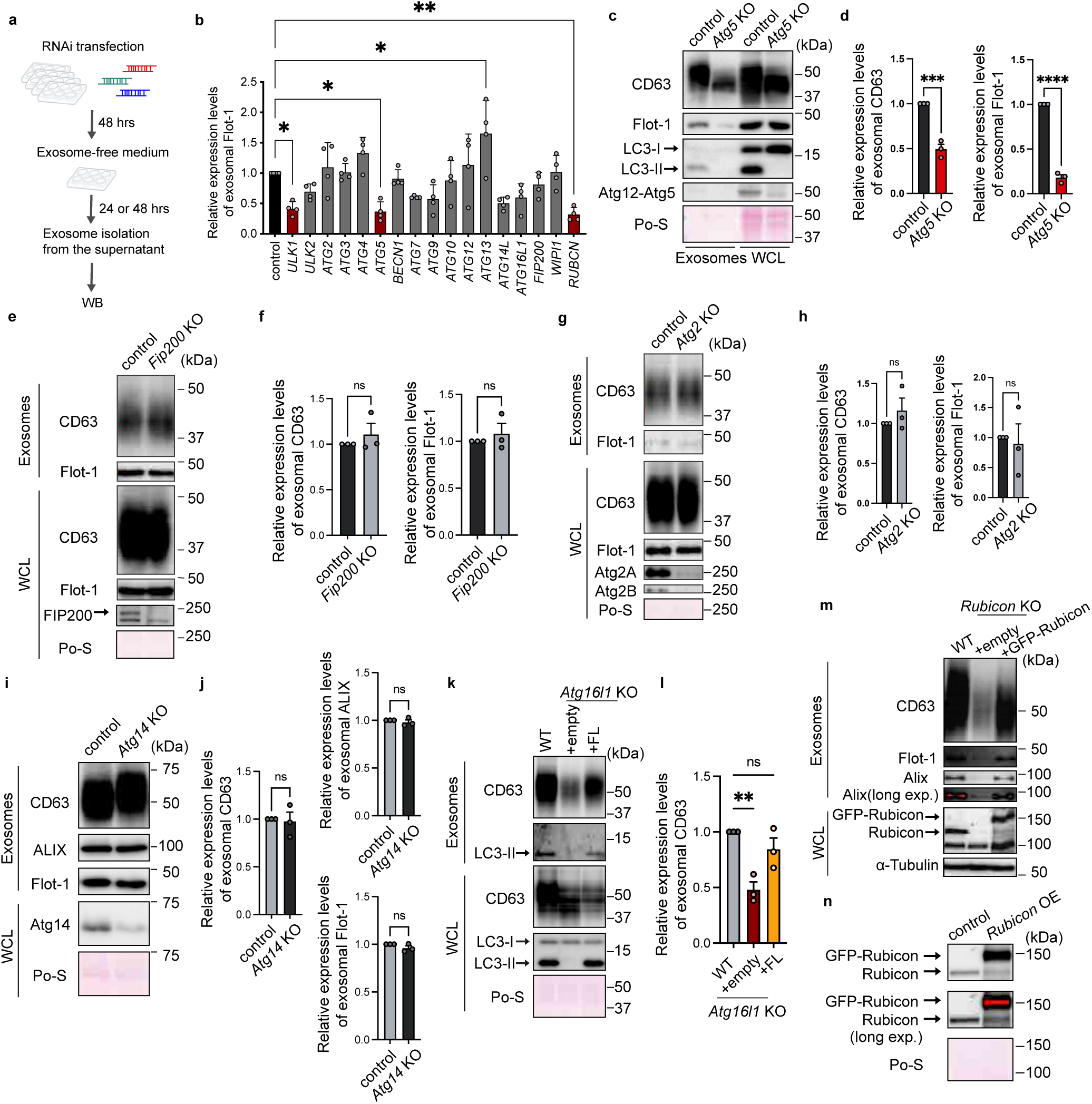
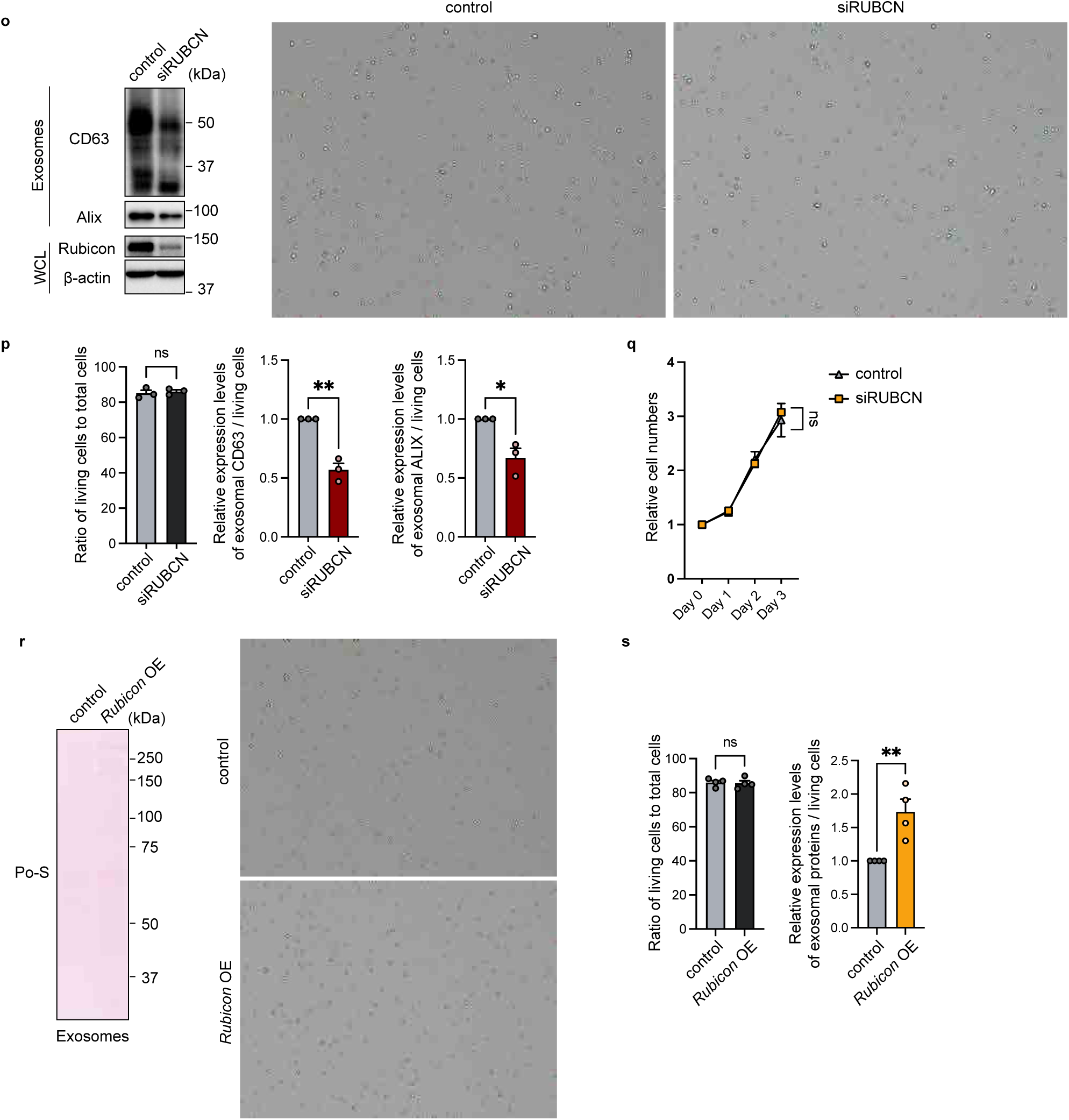
Rubicon promotes exosome biogenesis independently of autophagic pathways. (a) A schematic diagram of exosome isolation from culture medium of hMSCs transfected with siRNAs. (b) Quantification of exosomal Flot-1 levels in cells with knockdown of the genes indicated in Fig. 1a. n = 4. *p < 0.05, **p < 0.01 by one-way ANOVA followed by Dunnet’s test. (c) Immunoblotting of the indicated proteins in the exosome fractions and WCLs obtained from control and *Atg5*-KO MEFs. The loading amount for each sample was set as follows: half of the exosome sample collected from 1 ml of culture medium, and 6 µg for the WCL. (d) Quantification of exosomal levels of CD63 and Flot-1 in (d). Bars represent means ± SEM. n= 3 independent experiments. ***p < 0.001, ****p < 0.0001 by a two-tailed Student’s t-test. (e) Immunoblotting of the indicated proteins in the exosome fractions and WCLs obtained from control and *Fip200*-KO MEFs. The loading amount for each sample was set as follows: half of the exosome sample collected from 1 ml of culture medium, and 6 µg for the WCL. (f) Quantification of exosomal levels of CD63 and Flot-1 in (f). Bars represent means ± SEM. n = 3 independent experiments. ns, not significant by a two-tailed Student’s t-test. (g) Immunoblotting of the indicated proteins in the exosome fractions and WCLs obtained from control and *Atg2-KO* MEFs. The loading amount for each sample was set as follows: half of the exosome sample collected from 1 ml of culture medium, and 6 µg for the WCL. (h) Quantification of exosomal levels of CD63 and Flot-1 in (h). Bars represent means ± SEM. n = 3 independent experiments. ns, not significant by a two-tailed Student’s t-test. (i) Immunoblotting of the indicated proteins in the exosome fractions and WCLs obtained from control and *Atg14-KO* MEFs. The loading amount for each sample was set as follows: half of the exosome sample collected from 1 ml of culture medium, and 6 µg for the WCL. (j) Quantification of exosomal levels of CD63, ALIX and Flot-1 in (i). Bars represent means ± SEM. n = 3 independent experiments. ns, not significant by a two-tailed Student’s t-test. (k) Immunoblotting of the indicated proteins in the exosome fractions and WCLs obtained from WT or *Atg16l1*-KO MEFs expressing an empty plasmid (+empty) or ATG16L1 plasmid (+FL). n = 3. The loading amount for each sample was set as follows: half of the exosome sample collected from 1 ml of culture medium, and 6 µg for the WCL. (l) Quantification of exosomal levels of CD63 in (k). Bars represent means ± SEM. n = 3 independent experiments. ns, not significant; **p < 0.01 by one-way ANOVA followed by Dunnet’s test. (m) Immunoblotting of the indicated proteins in the exosome fractions and WCLs obtained from WT MEFs and *Rubicon*-KO MEFs expressing an empty plasmid (+empty) or a GFP-Rubicon plasmid (+GFP-Rubicon). The loading amount for each sample was set as follows: half of the exosome sample collected from 1 ml of culture medium, and 6 µg for the WCL. (n) Immunoblotting of the indicated proteins in the WCLs obtained from control and *Rubicon*-OE MEFs. The loading amount for each sample was set as 10 µg. (o) Left, immunoblotting of the indicated proteins in the exosome fractions and WCLs obtained from control and *RUBCN*-knockdown hMSCs. Right, representative image of Trypan Blue staining, live cells are marked in green and dead cells in red, in indicated cells. The loading amount for each sample was set as follows: half of the exosome sample collected from 1 ml of culture medium, and 6 µg for the WCL. (p) Left, quantification of the ratio of live cells to total cells in (o). Right, quantification of exosomal levels of CD63 and ALIX normalized by the number of live cells in (o). Bars represent means ± SEM. n = 3 independent experiments. ns, not significant; *p < 0.05, **p < 0.01 by a two-tailed Student’s t-test. (q) Relative numbers of total cells in control and *RUBCN* knockdown hMSCs. Bars represent means ± SEM. n = 3 independent experiments. ns, not significant by a two-tailed Student’s t-test. (r) Left, ponceau S staining in the exosome fractions obtained from control and *Rubicon-* overexpressing MEFs. Right, representative image of Trypan Blue staining, live cells are marked in green and dead cells in red, in indicated cells. The loading amount for each sample was set as follows: half of the exosome sample collected from 1 ml of culture medium. (s) Left, quantification of the ratio of live cells to total cells in (r). Right, quantification of exosomal levels of ponceau S staining normalized by the number of live cells in (r). Bars represent means ± SEM. n = 4 independent experiments. ns, not significant; **p < 0.01 by a two-tailed Student’s t-test.

**Extended Data Fig. 2.**
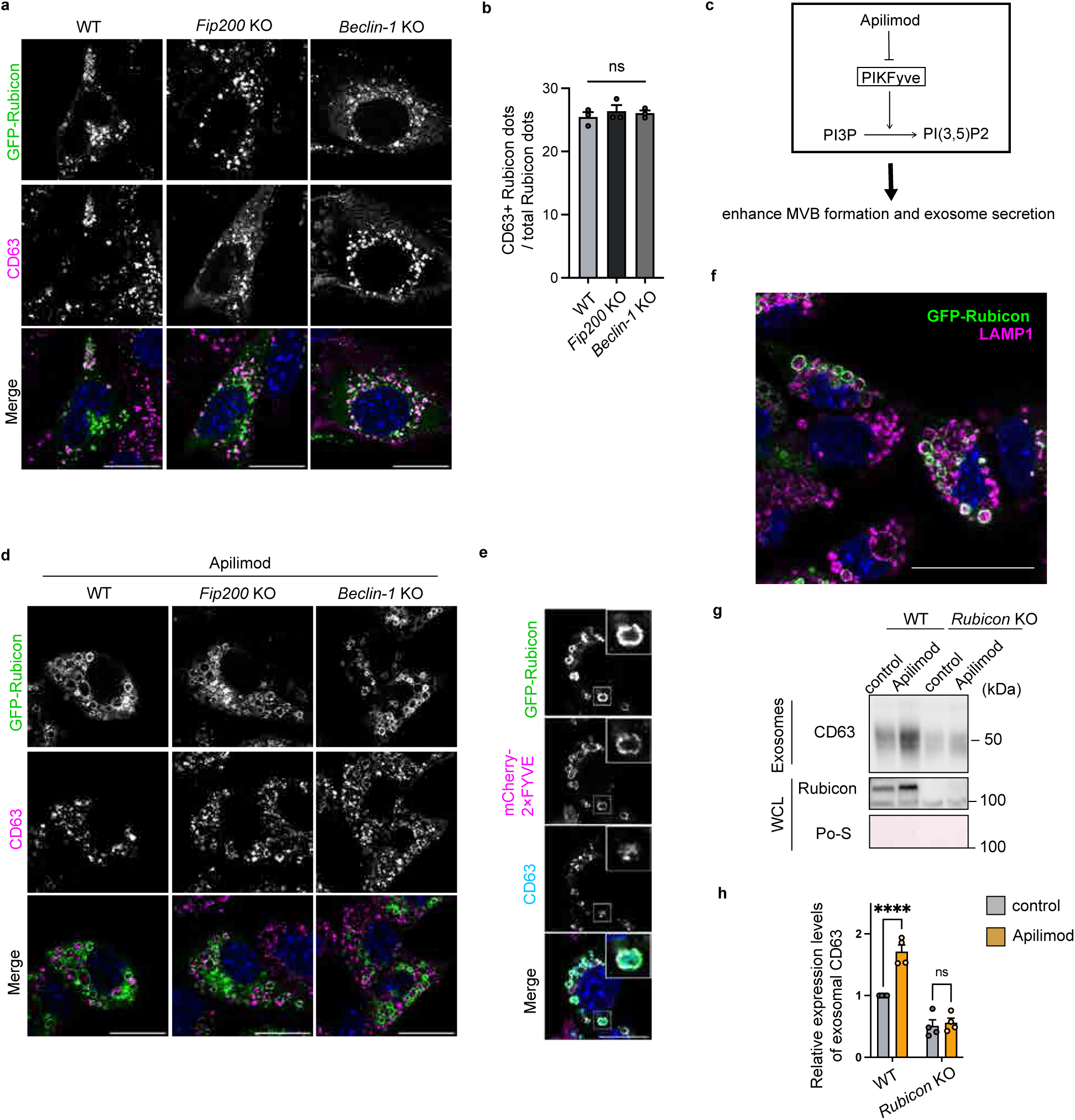
Apilimod treatment promotes exosome biogenesis in a Rubicon-dependent manner. (a) Immunofluorescent images of CD63 (magenta) and DAPI (blue) in WT, *Fip200*-KO, and *Beclin-1*-KO MEFs expressing GFP-Rubicon. Scale bars, 50 μm. (b) Quantification of the ratio of CD63- and Rubicon-positive dots relative to Rubicon-positive dots in indicated cells. n = 3. ns, not significant by one-way ANOVA followed by Dunnet’s test. (c) A schematic diagram of pharmacological inhibition of PIKFyve by Apilimod. (d) Immunofluorescent images of CD63 (magenta) and DAPI (blue) in WT, *Fip200*-KO, and *Beclin-1*-KO MEFs expressing GFP-Rubicon with 0.5 μM Apilimod treatment for 1 hour. Scale bars, 50 μm. (e) Immunofluorescent images of CD63 (cyan) and DAPI (blue) in MEFs expressing GFP-Rubicon and mCherry-2×FYVE with Apilimod treatment. Scale bars, 50 μm. (f) Immunofluorescent images of LAMP1 (magenta) and DAPI (blue) in MEFs expressing GFP-CD63 with Apilimod treatment. Scale bars, 50 μm. (g) Immunoblotting of the indicated proteins in the exosome fractions and WCLs obtained from WT and *Rubicon*-KO MEFs with or without Apilimod treatment. The loading amount for each sample was set as follows: half of the exosome sample collected from 1 ml of culture medium, and 6 µg for the WCL. (h) Quantification of exosomal CD63 levels in (g). Bars represent means ± SEM. n = 4 independent experiments. ns, not significant; ****p < 0.0001 by two-way ANOVA.

**Extended Data Fig. 3.**
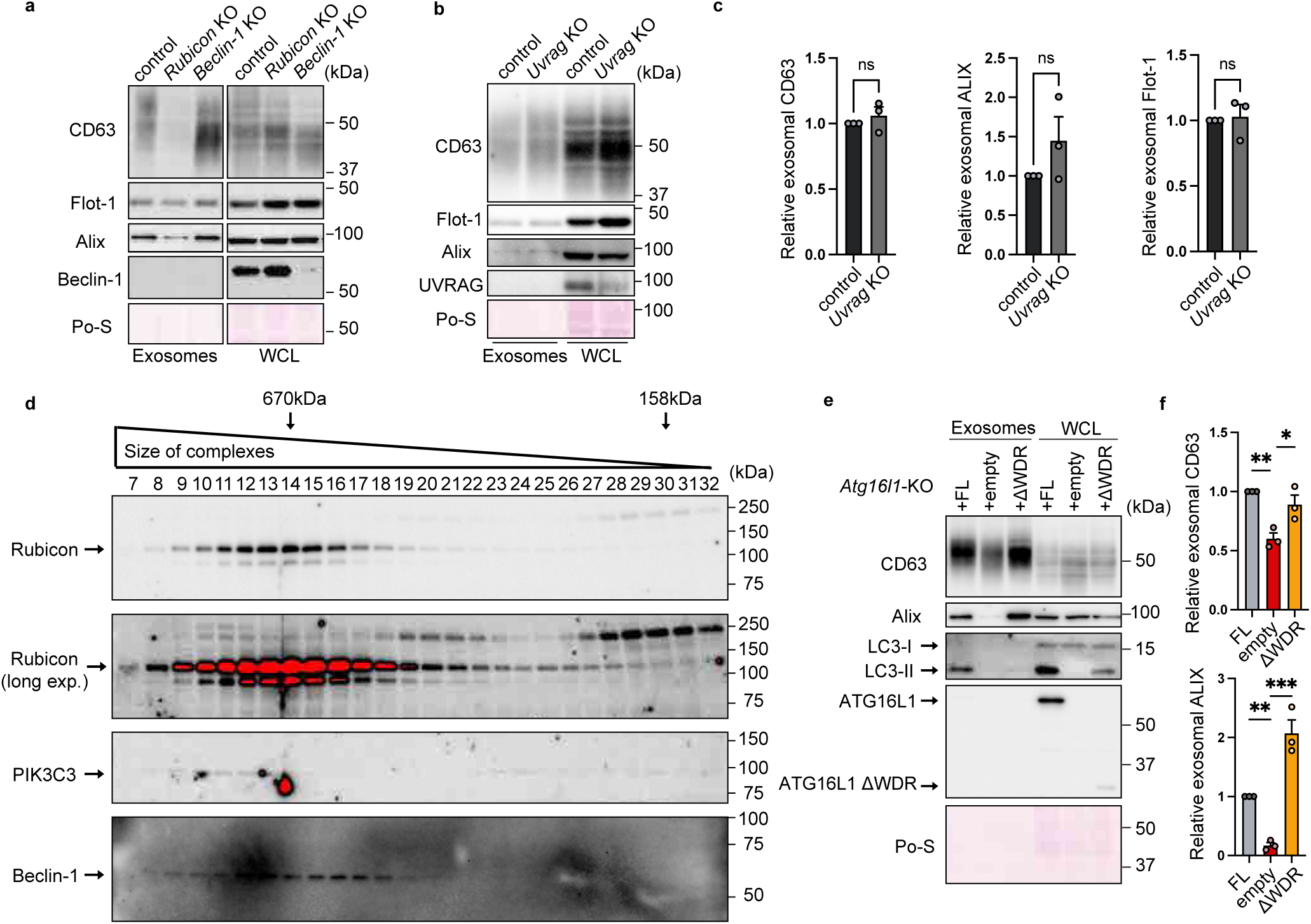
Both canonical and non-canonical autophagic pathways are dispensable for exosome secretion. (a) Immunoblotting of the indicated proteins in the exosome fractions and WCLs obtained from control, *Rubicon*-KO, and *Beclin-1-KO* MEFs. The loading amount for each sample was set as follows: half of the exosome sample collected from 1 ml of culture medium, and 6 µg for the WCL. (b) Immunoblotting of the indicated proteins in the exosome fractions and WCLs obtained from control and *Uvrag-KO* MEFs. The loading amount for each sample was set as follows: half of the exosome sample collected from 1 ml of culture medium, and 10 µg for the WCL. (c) Quantification of exosomal levels of CD63, Flot-1, and ALIX in (b). Bars represent means ± SEM. n = 3 independent experiments. ns, not significant by a two-tailed Student’s t-test. (d) Immunoblotting of the indicated proteins in gel filtration fractions obtained from MSCs. (e) Immunoblotting of the indicated proteins in the exosome fractions and WCLs obtained from *Atg16l1*-KO MEFs expressing an ATG16L1 plasmid (+FL), an empty plasmid (+empty), or a WD-repeat domain–deleted ATG16L1 plasmid (+ΔWDR). The loading amount for each sample was set as follows: half of the exosome sample collected from 1 ml of culture medium, and 6 µg for the WCL. (f) Quantification of exosomal levels of CD63 and ALIX in (d). Bars represent means ± SEM. n = 3. *p < 0.05, **p < 0.01, ***p < 0.001 by one-way ANOVA followed by Dunnett’s test.

**Extended Data Fig. 4.**
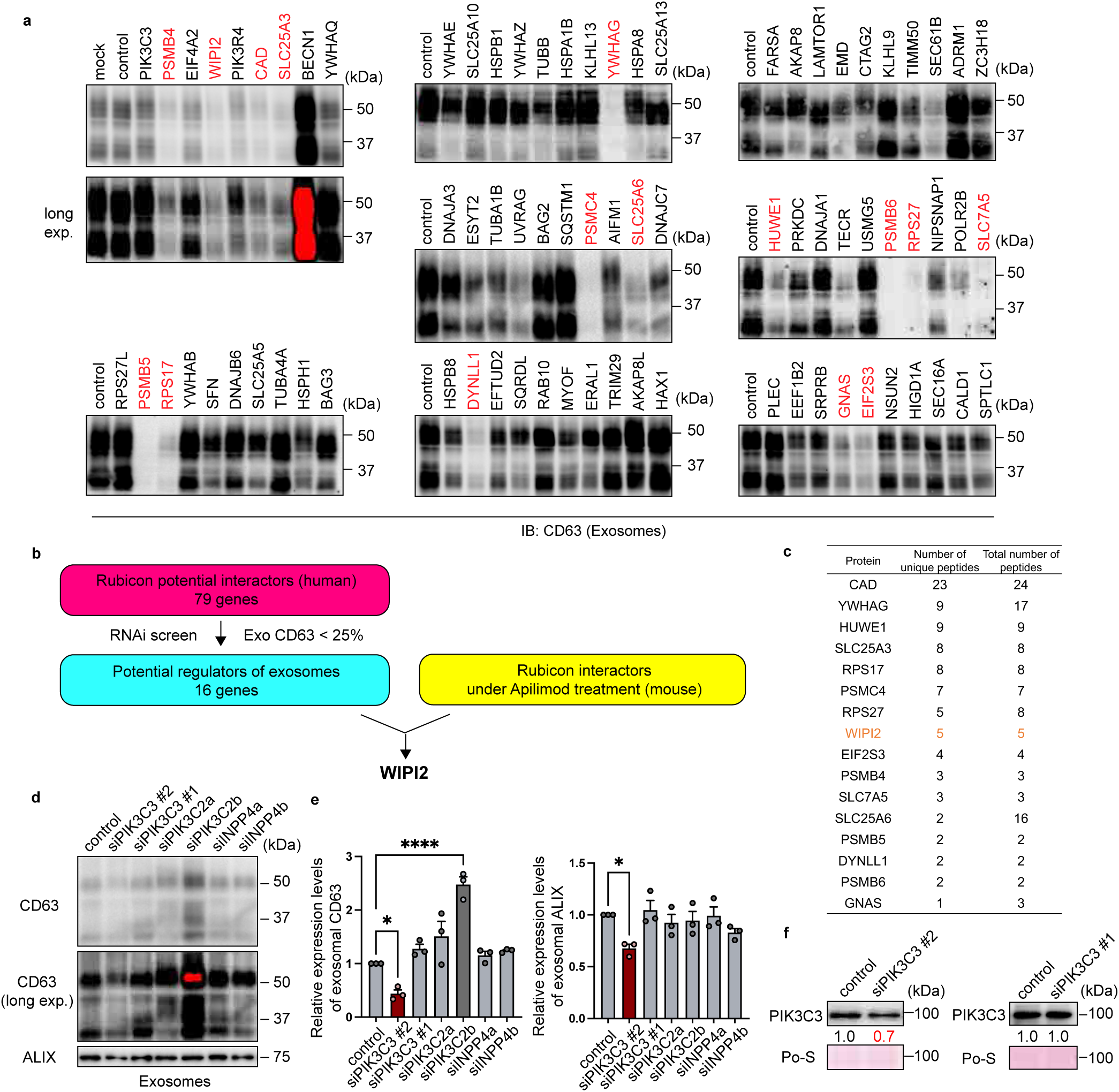
RNAi screening of exosome regulators identifies multiple candidates from Rubicon-interacting proteins. (a) Immunoblotting of CD63 in the exosome fractions isolated from culture medium of hMSCs with knockdown of the indicated genes. The loading amount for each sample was set as follows: half of the exosome sample collected from 1 ml of culture medium. (b) A schematic diagram showing identification of WIPI2 by the interactome analysis and RNAi screen of exosome secretions. (c) The number of unique peptides and total peptides that are potential regulators of exosomes in (b), as detected in Rubicon-interactome analysis using HeLa Kyoto cells. (d) Immunoblotting of CD63 and ALIX in the exosome fractions isolated from the culture medium of hMSCs with knockdown of the indicated genes. The loading amount for each sample was set as follows: half of the exosome sample collected from 1 ml of culture medium. (e) Quantification of exosomal levels of CD63 and ALIX in (d). Bars represent means ± SEM. n = 3 independent experiments. *p < 0.05, ****p < 0.0001 by one-way ANOVA followed by Dunnett’s test. (f) Immunoblotting of indicated proteins in hMSCs with knockdown of the indicated genes. The loading amount for each sample was set as 10 µg.

**Extended Data Fig. 5.**
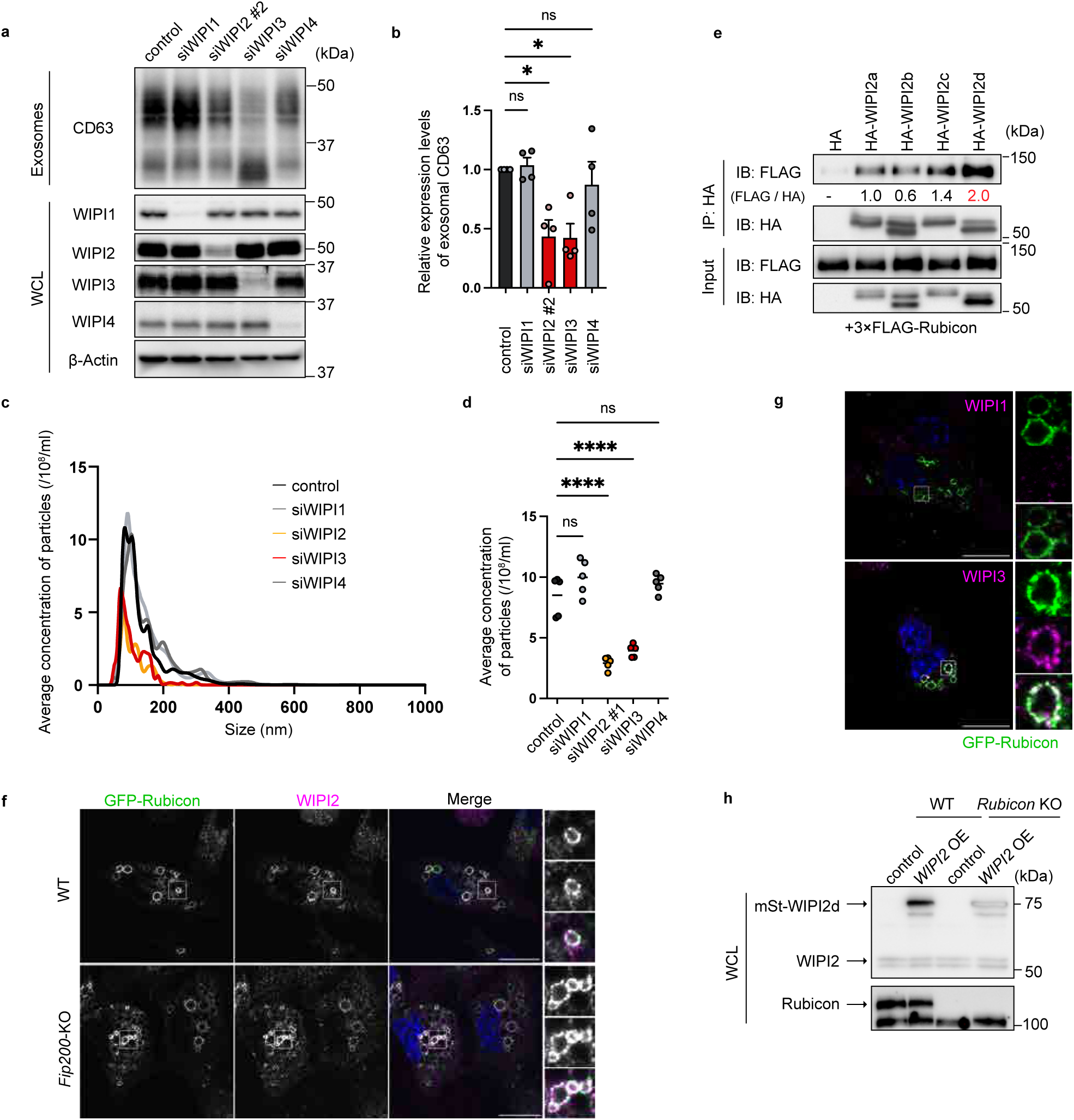
Rubicon recruits WIPI proteins to endosomes, which is necessary for exosome biogenesis. (a) Immunoblotting of the indicated proteins in the exosome fractions and WCLs obtained from hMSCs with knockdown of the indicated genes. The loading amount for each sample was set as follows: half of the exosome sample collected from 1 ml of culture medium, and 6 µg for the WCL. (b) Quantification of exosomal levels of CD63 in (a). Bars represent means ± SEM. n = 4 independent experiments. ns, not significant; *p < 0.05 by one-way ANOVA followed by Dunnett’s test. (c) NTA of EVs purified from the cultured medium of hMSCs with knockdown of indicated genes using ultracentrifugation. (d) Quantification of the total particle concentration of the EVs in (c). Bars represent means. n = 5 independent experiments. ns, not significant; ****p < 0.0001 by one-way ANOVA followed by Dunnett’s test. (e) Immunoprecipitation assay. HEK293T cells were transfected with the indicated plasmids for 24 hours, and then the cells were lysed and immunoprecipitated with anti-HA antibody. Precipitates were subjected to immunoblotting with the indicated antibodies. (f) Immunofluorescent images of WIPI2 (magenta) and DAPI (blue) in WT MEFs or *Fip200*-KO MEFs expressing GFP-Rubicon with Apilimod treatment. Scale bars, 50 μm. (g) Immunofluorescence images of WIPI1 or WIPI3 (magenta) and DAPI (blue) in MEFs expressing GFP-Rubicon with Apilimod treatment. Scale bars, 50 μm. (h) Immunoblotting of indicated proteins in the WCLs obtained from WT and *Rubicon*-KO MEFs expressing an empty plasmid (control) or an mStrawberry-WIPI2d plasmid (WIPI2 OE). The loading amount for each sample was set as 6 µg.

**Extended Data Fig. 6.**
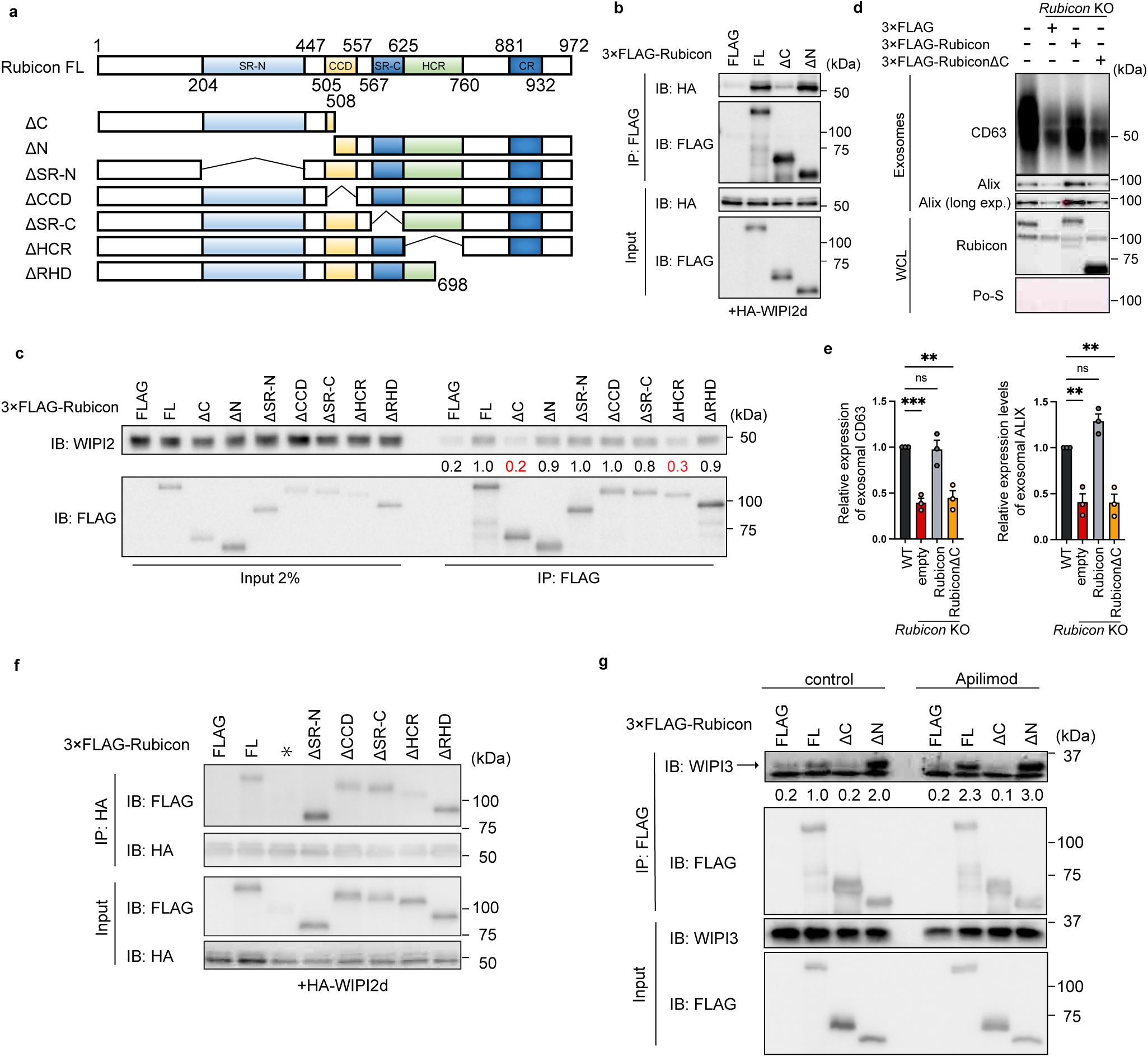
The helix-coil-rich region in Rubicon is necessary for its interaction with WIPI2. (a) Schematic representation of Rubicon and its truncated mutants. Rubicon FL, 3×FLAG-Rubicon (1–972); ΔC, 3×FLAG-Rubicon (1–508); ΔN, Rubicon (509–972); ΔSR-N, 3×FLAG-Rubicon (1–203, 448–972); ΔCCD, 3×FLAG-Rubicon (1–504, 558–972); ΔSR-C, 3×FLAG-Rubicon (1–566, 626–972); ΔHCR, 3×FLAG-Rubicon (1-625, 761–972); ΔRHD, 3×FLAG-Rubicon (1–698). (b) Immunoprecipitation assay. HEK293T cells were transfected with HA-WIPI2d and 3×FLAG (FLAG) or the Rubicon mutant plasmids for 24 hours, and then the cells were lysed and immunoprecipitated with anti-FLAG antibody. Precipitates were subjected to immunoblotting with the indicated antibodies. (c) Immunoprecipitation assay. HEK293T cells were transfected with the indicated plasmids for 24 hours, and then the cells were lysed and immunoprecipitated with anti-FLAG antibody. Precipitates were subjected to immunoblotting with the indicated antibodies. (d) Immunoblotting of the indicated proteins in the exosome fractions and WCLs obtained from WT and *Rubicon*-KO MEFs expressing the indicated plasmids. The loading amount for each sample was set as follows: half of the exosome sample collected from 1 ml of culture medium, and 6 µg for the WCL. (e) Quantification of exosomal levels of CD63 and ALIX in (d). Bars represent means. n = 3. ns, not significant; **p < 0.01, ***p < 0.001 by one-way ANOVA followed by Dunnett’s test. (f) Immunoprecipitation assay. HEK293T cells were transfected with HA-WIPI2d and the Rubicon mutant plasmids (* represents missing transfection) for 24 hours, and then the cells were lysed and immunoprecipitated with anti-HA antibody. Precipitates were subjected to immunoblotting with the indicated antibodies. (g) Immunoprecipitation assay. HEK293T cells were transfected with the indicated plasmids for 24 hours with or without Apilimod treatment, and then the cells were lysed and immunoprecipitated with anti-FLAG antibody. Precipitates were subjected to immunoblotting with the indicated antibodies.

**Extended Data Fig. 7.**
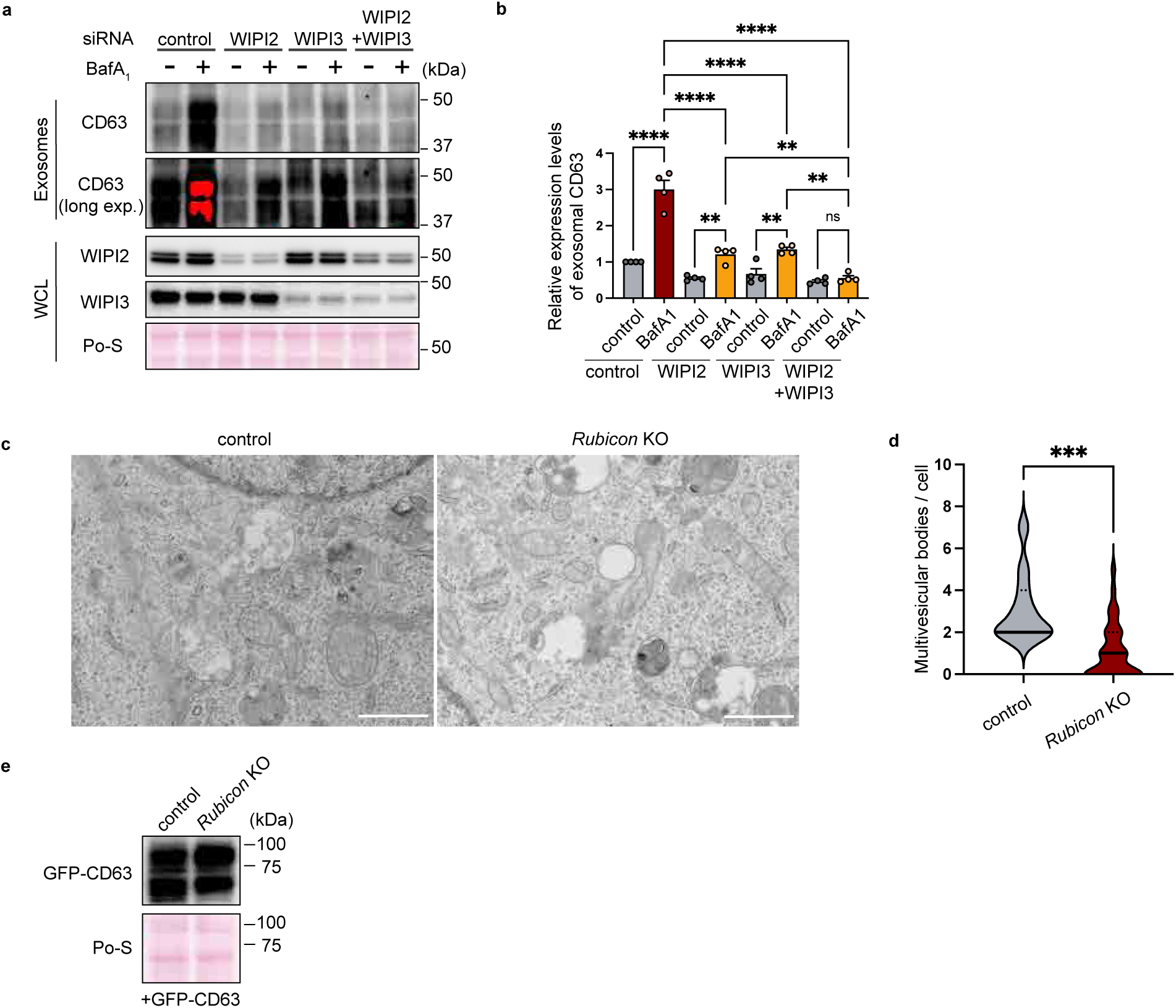
The Rubicon-WIPI axis is required for MVB formation. (a) Immunoblotting of the indicated proteins in the exosome fractions and WCLs obtained from hMSCs treated with or without bafilomycin A1 for 3 hours and with knockdown of the indicated genes. The loading amount for each sample was set as follows: half of the exosome sample collected from 1 ml of culture medium, and 6 µg for the WCL. (b) Quantification of exosomal levels of CD63 in (a). Bars represent means ± SEM. n = 4 independent experiments. ns, not significant; **p < 0.01, ****p < 0.0001 by one-way ANOVA followed by Tukey’s test. (c) Representative electron microscopic images of multivesicular bodies (MVBs) in control and *Rubicon*-KO MEFs. (d) Violin plots showing the number of MVBs per cell in (c). WT, n = 11; *Rubicon* KO, n = 35. The solid line denotes the median, and dotted lines define the quartiles; ***p < 0.001 by a two-tailed Student’s t-test. (e) Immunoblotting of GFP-CD63 in the WCLs obtained from control or *Rubicon*-KO MEFs stably expressing GFP-CD63. The loading amount for each sample was set as 6 µg.

**Extended Data Fig. 8.**
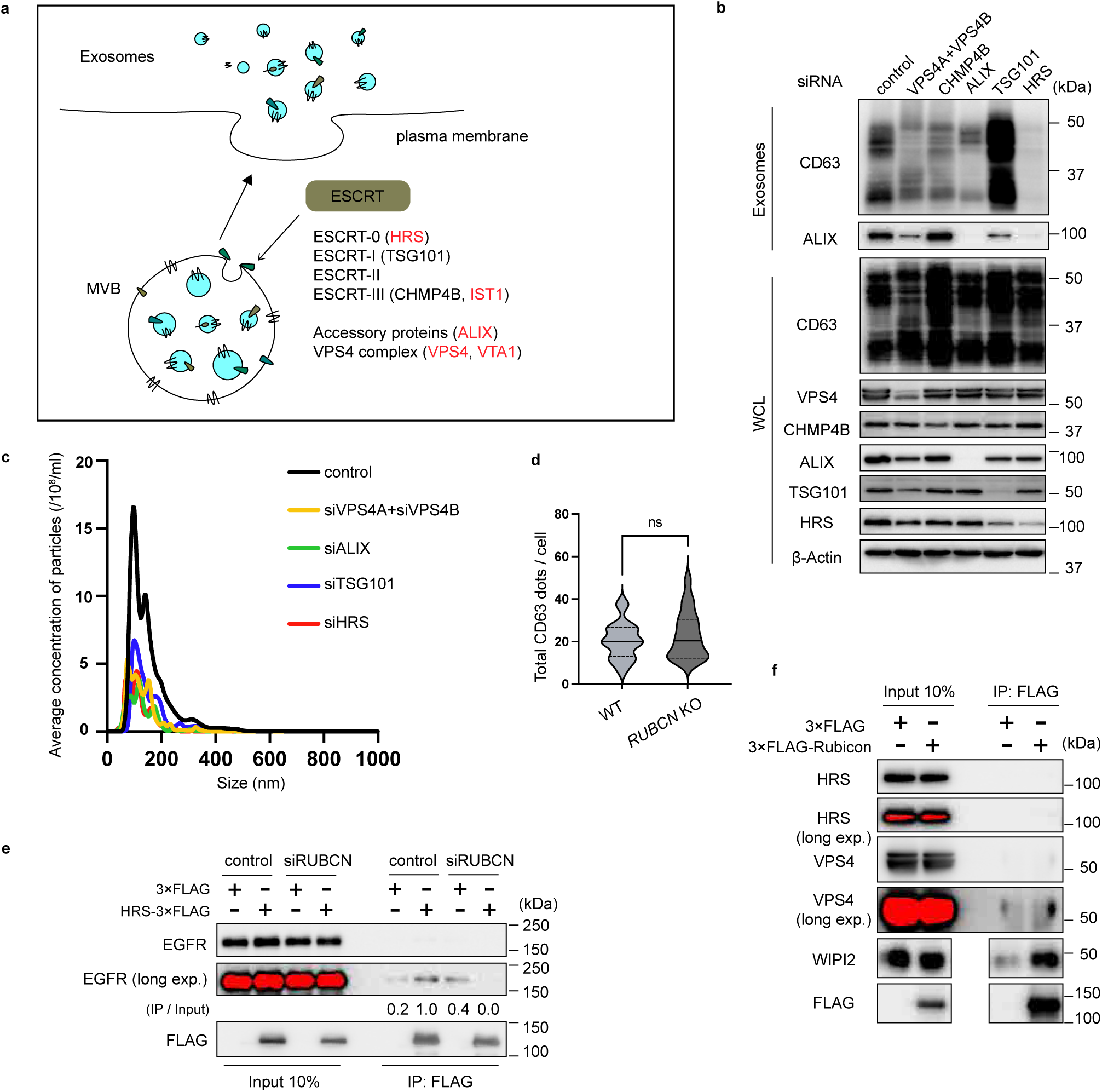
The ESCRT machinery interacts with WIPI2 and is necessary for exosome biogenesis. (a) Illustration of the ESCRT machinery and MVB formation process. Red letters represent ESCRT components and associated proteins that demonstrate interaction with WIPI2d in Fig. 4a. (b) Immunoblotting of the indicated proteins in the exosome fractions and WCLs obtained from hMSCs with knockdown of the indicated genes. The loading amount for each sample was set as follows: half of the exosome sample collected from 1 ml of culture medium, and 6 µg for the WCL. (c) NTA of EVs purified from the cultured medium of hMSCs with knockdown of indicated genes using ultracentrifugation. (d) Violin plot showing the total number of CD63 dots in the indicated cells in Fig. 4c. n = 21. The solid line denotes the median, and the dotted lines define the quartiles. ns, not significant by a two-tailed Student’s t-test. (e) Immunoprecipitation assay. HEK293T cells were knocked down with the indicated genes and transfected with the indicated plasmids, and then the cells were lysed and immunoprecipitated with anti-FLAG antibody. Precipitates were subjected to immunoblotting with the indicated antibodies. (f) Immunoprecipitation assay. HEK293T cells were transfected with the indicated plasmids, and then the cells were lysed and immunoprecipitated with anti-FLAG antibody. Precipitates were subjected to immunoblotting with the indicated antibodies.

**Extended Data Fig. 9.**
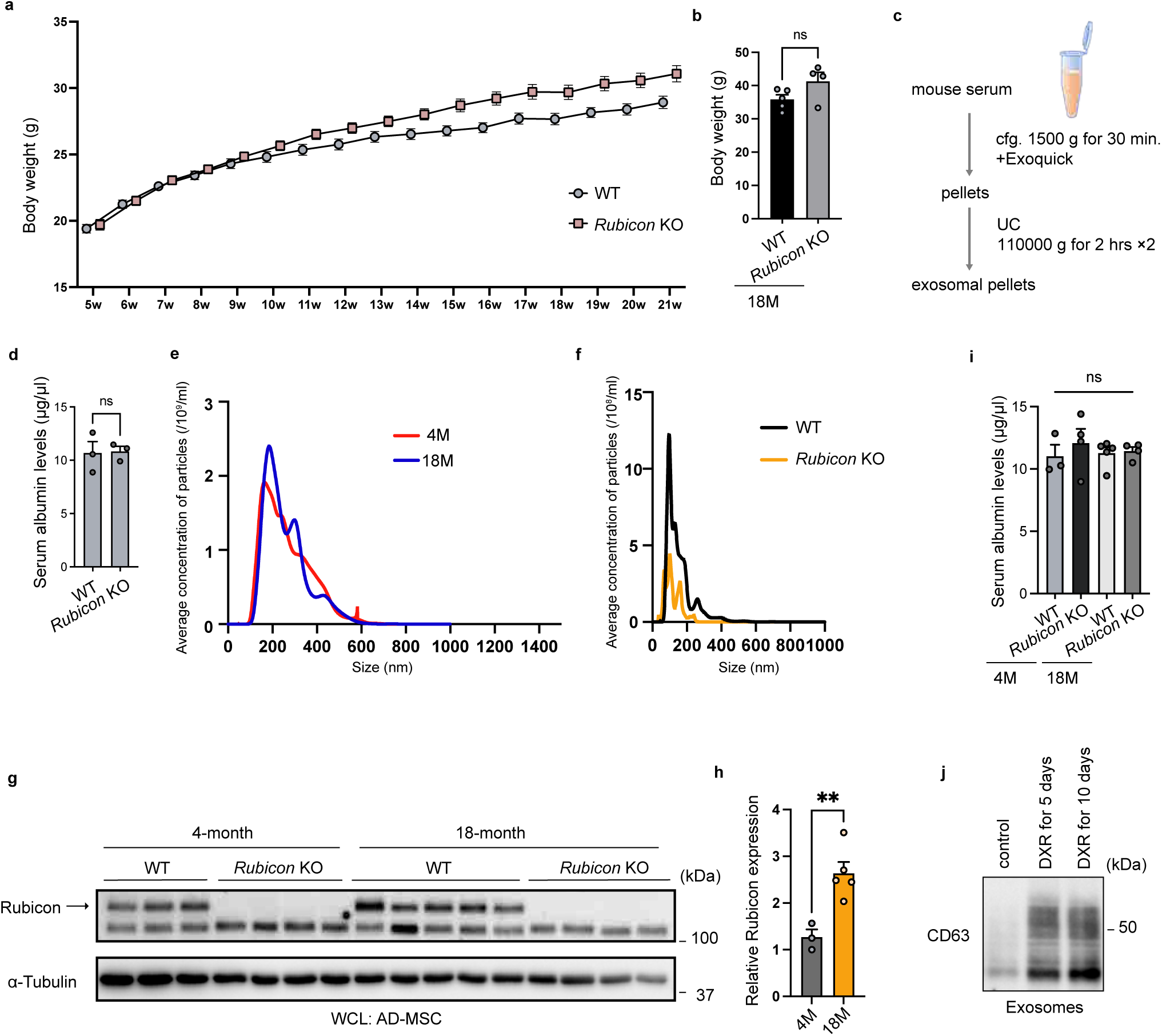
*Rubicon* knockout suppresses the age-related increase in exosome secretion *in vivo*. (a) Body weight chart for mice of the indicated genotypes on a normal chow diet. WT, n = 21; *Rubicon* KO, n = 21. Bars represent means ± SEM. (b) Quantification of body weight of 18-month-old mice of the indicated genotypes. WT, n = 5; *Rubicon*-KO, n = 4. Bars represent means ± SEM. (c) A schematic diagram of serum exosome isolation. (d) Serum albumin levels obtained from control or *Rubicon*-KO mice. n = 3. Bars represent means ± SEM. ns, not significant; by a two-tailed Student’s t-test. (e) NTA of large EVs purified from serum obtained from 4- or 18-month-old mice using a PS-affinity kit. (f) NTA of EVs purified from primary AD-MSCs obtained from control or *Rubicon*-KO mice using ultracentrifugation. (g) Immunoblotting of the indicated proteins in the WCLs of primary AD-MSCs obtained from young or aged mice of the indicated genotypes. 4-month-old WT, n = 3; 4-month-old *Rubicon*-KO, n = 4; 18-month-old WT, n = 5; 18-month-old *Rubicon* KO, n = 4. The loading amount for each sample was set as 6 µg. (h) Quantification of Rubicon levels normalized by α-tubulin in (c). Bars represent means ± SEM. 4 months old, n = 3; 18 months old, n = 5; **p < 0.01 by a two-tailed Student’s t-test. (i) Serum albumin levels obtained from indicated mice. 4-month-old WT, n = 3; 4-month-old Rubicon-KO, n = 4; 18-month-old WT, n = 5; 18-month-old Rubicon KO, n = 4. Bars represent means ± SEM. ns, not significant; by one-way ANOVA followed by Tukey’s test. (j) Immunoblotting of the indicated proteins in the exosome fractions obtained from RPE-1 cells with the indicated DXR treatment. The loading amount for each sample was set as follows: half of the exosome sample collected from 1 ml of culture medium.

**Extended Data Fig. 10.**
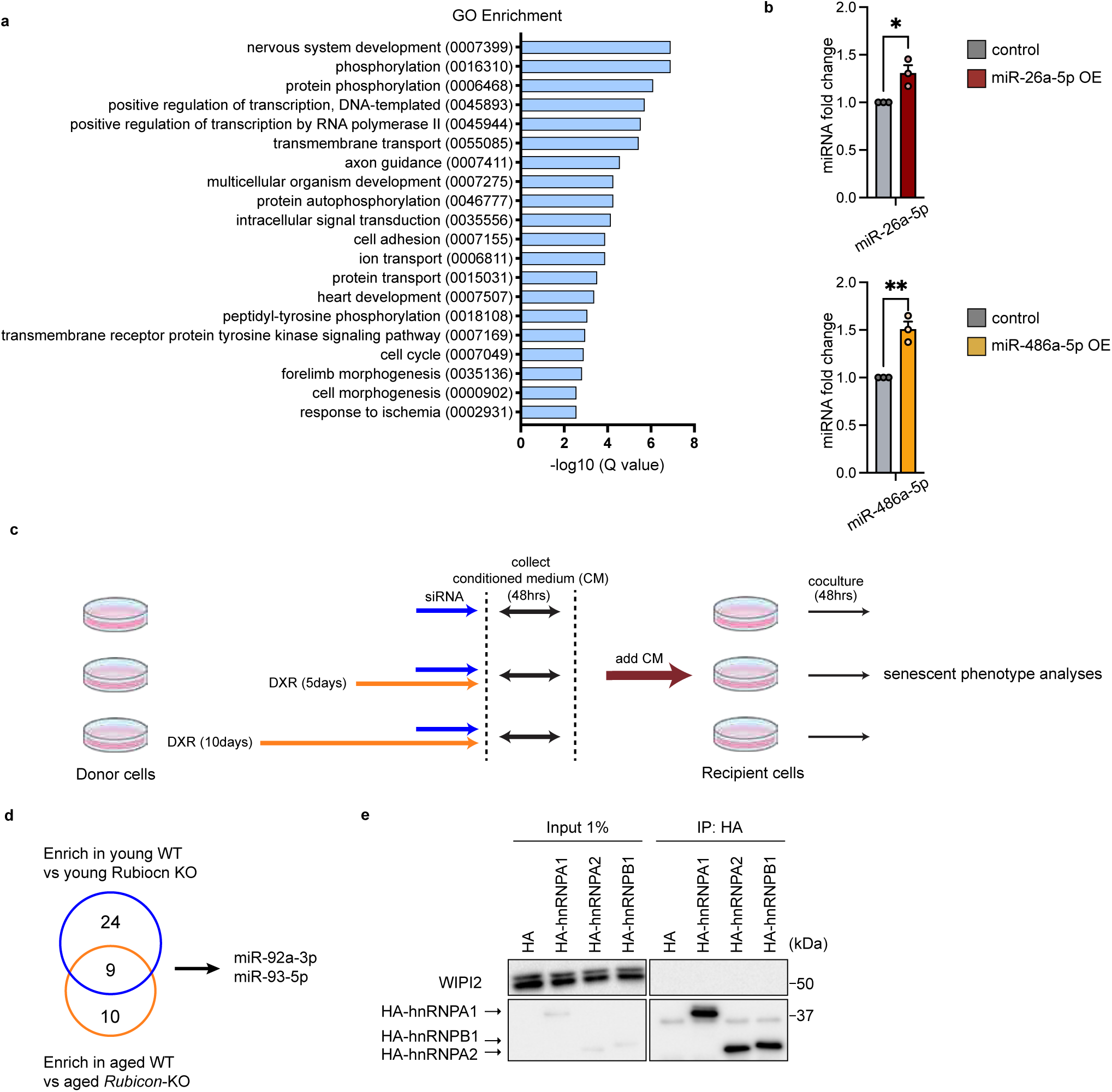
Rubicon is essential for the age-dependent alteration of the exosomal microRNA profile that exacerbate cellular senescence. (a) Gene ontology enrichment analysis of the target genes of the 10 miRNAs enriched in the serum EVs in Fig. 6b. X-axis, −log_10_ q-value of duplicate (n = 3) experiments. (b) Fold change of indicated miRNAs levels in indicated cells. n = 3. *p < 0.05, **p < 0.01 by a two-tailed Student’s t-test. (c) A schematic diagram of treatment with CM of senescent donor cells to recipient cells. (d) Venn diagrams showing the numbers of DEGs in serum EVs from young WT relative to young *Rubicon*-KO (blue) and age WT relative to aged *Rubicon*-KO (yellow) mice.

**Extended Data Table. 1 Interactome analysis of Rubicon using HeLa Kyoto cells**

**Extended Data Table. 2 Interactome analysis of Rubicon using MEFs under Apilimod treatment**

**Extended Data Table. 3 Interactome analysis of WIPI2d**

**Extended Data Table. 4 The siRNAs used in this study**

